# Cargo-Adaptor Cooperation Programs Retromer Coat Architecture

**DOI:** 10.64898/2026.05.05.722683

**Authors:** Marta Pardo-Piñón, Raffaele Coray, Pengwei Zhang, Changin Oh, Adriana L Rojas, Daniel DiMaio, Daniel Castaño-Díez, Aitor Hierro

## Abstract

Retromer drives endosomal cargo retrieval in combination with sorting nexin (SNX) adaptors, but how adaptor-cargo combinations specify coat architecture remains unclear. We identify sorting nexin 12 (SNX12) as the retromer adaptor required for human papillomavirus 16 (HPV16) infection and show that the viral L2 capsid protein tail directly engages SNX12-retromer complexes to trigger membrane tubulation. The crystal structure reveals a conserved cargo-recognition mode, whereas cryo-electron tomography of reconstituted assemblies shows retromer arches organized into two lattice configurations stabilized by membrane-proximal interfaces. These lattices assemble as multi-start helices and accommodate curvature through hinge-like motions between arches. These findings establish cargo and adaptor identity as co-determinants of retromer coat architecture, revealing retromer as a programmable system capable of generating route-specific transport carriers.

---

Endosomal sorting contributes to cellular homeostasis determining whether membrane proteins are recycled or degraded. A central component of this system is the retromer complex, which retrieves selected cargo from endosomes and packages it into tubular carriers destined for the trans-Golgi network or the plasma membrane^1,2^. In metazoans, retromer is composed of a trimeric core of VPS26A/B, VPS29A/B/C, and VPS35 subunits^2–4^, working in concert with a subset of sorting nexins (SNXs) that serve as adaptors to direct cargo to the trans-Golgi network or plasma membrane^5,6^. Retromer dysfunction is linked to Alzheimer’s and Parkinson’s disease, metabolic disorders, and the intracellular replication of diverse pathogens^7–10^ underscoring the importance of understanding how this coat assembles and functions.

Retromer associates with SNX1/2-SNX5/6 heterodimers or with SNX3 monomers to facilitate endosome-to-Golgi retrieval via tubular endosomal carriers, while SNX27 guides endosome-to-plasma membrane recycling^11–15^. SNX3 and its close homologue SNX12 are composed of only a PX domain^16^. SNX3 cooperates with the retromer subunit VPS26 to directly bind cargo containing a Φx[LMV] motif, where Φ represents hydrophobic amino acids^17,18^. Consistent with this adaptor-cargo cooperation, yeast reconstitution studies indicate that cargo enhances retromer-dependent membrane remodeling^19^. Comparisons between yeast and mammalian assemblies, and between SNX-BAR and SNX-PX adaptors^20,21^, suggest a modular system where the core trimer and various adaptors jointly shape membrane curvature, but the structural principles underlying the contribution of cargo to coat geometry, have not been defined.

Human papillomaviruses (HPVs) are the causative agents of cervical and several other cancers, together accounting for approximately 5% of all human malignancies. To establish infection, HPV16 must navigate the endolysosomal system and deliver its genome to the nucleus. After cellular uptake, the viral capsid partially disassembles in the endosomal lumen, allowing the C-terminal tail of the minor capsid protein L2 to penetrate the endosomal membrane and protrude into the cytosol^22^. This exposed cytoplasmic tail engages multiple host trafficking factors including retromer^22–24^. Two short retromer-binding motifs in L2, ^446^FYL^448^ and ^452^YYML^455^, are among the first regions exposed to the cytosol and are essential for viral exit from endosomes and transport to the trans-Golgi network (TGN). Mutation of these motifs, or depletion of retromer, causes HPV to accumulate in endosomes and prevents trafficking to the TGN, thereby blocking infection^23,25^. How L2 recruits retromer, which SNX adaptor is required, and how this engagement drives the assembly of a tubular transport carrier, have all remained open questions.

Here we identify SNX12 as the retromer adaptor selectively required for HPV16 endosomal escape. Using purified components, we reconstitute cargo-dependent membrane tubulation and determine the crystal structure of the SNX12-retromer-L2 complex, revealing the molecular basis of viral cargo recognition. Cryo-electron tomography of reconstituted assemblies further shows that retromer arches organize into pseudohelical lattices stabilized by membrane-proximal contacts that align cargo-binding sites across neighboring complexes. Together, these results demonstrate that adaptor-cargo combinations program retromer coat architecture, establishing cargo as an integral structural component of the coat.

## Results

### SNX12 is important for HPV16 Infection

To determine SNX proteins involved in HPV infection, we generated clonal SNX3, SNX12, and SNX3/12 double knockout (KO) HeLa cells. The extinction of SNX3 and SNX12 was demonstrated by Western blotting (Fig. 1a), which also showed that KO of SNX12 did not affect expression of SNX3, and vice versa. To assess the effect of SNX3 and SNX12 KO on HPV infection, these KO cells were infected with HPV16 pseudovirus (PsV), which consists of a capsid comprised of L1 and L2 containing a reporter plasmid expressing HcRed. Infectivity was assessed at 48 hours post-infection (hpi) by using flow cytometry to measure HcRed fluorescence in infected cells. Although there was considerably clone-to-clone variability, the deletion of SNX12 inhibited HPV infectivity compared to control cells on average by approximately 85%, while SNX3 deletion inhibited infection by less than 35% (Fig. 1b). The SNX3/SNX12 double knockout cells also exhibited a significant reduction in HPV PsV infectivity, similar to the effect of deleting SNX12 alone. The different infectivity of different SNX12 KO clones may be due to compensatory changes occurring during the derivation of the cells.

**Fig. 1:**
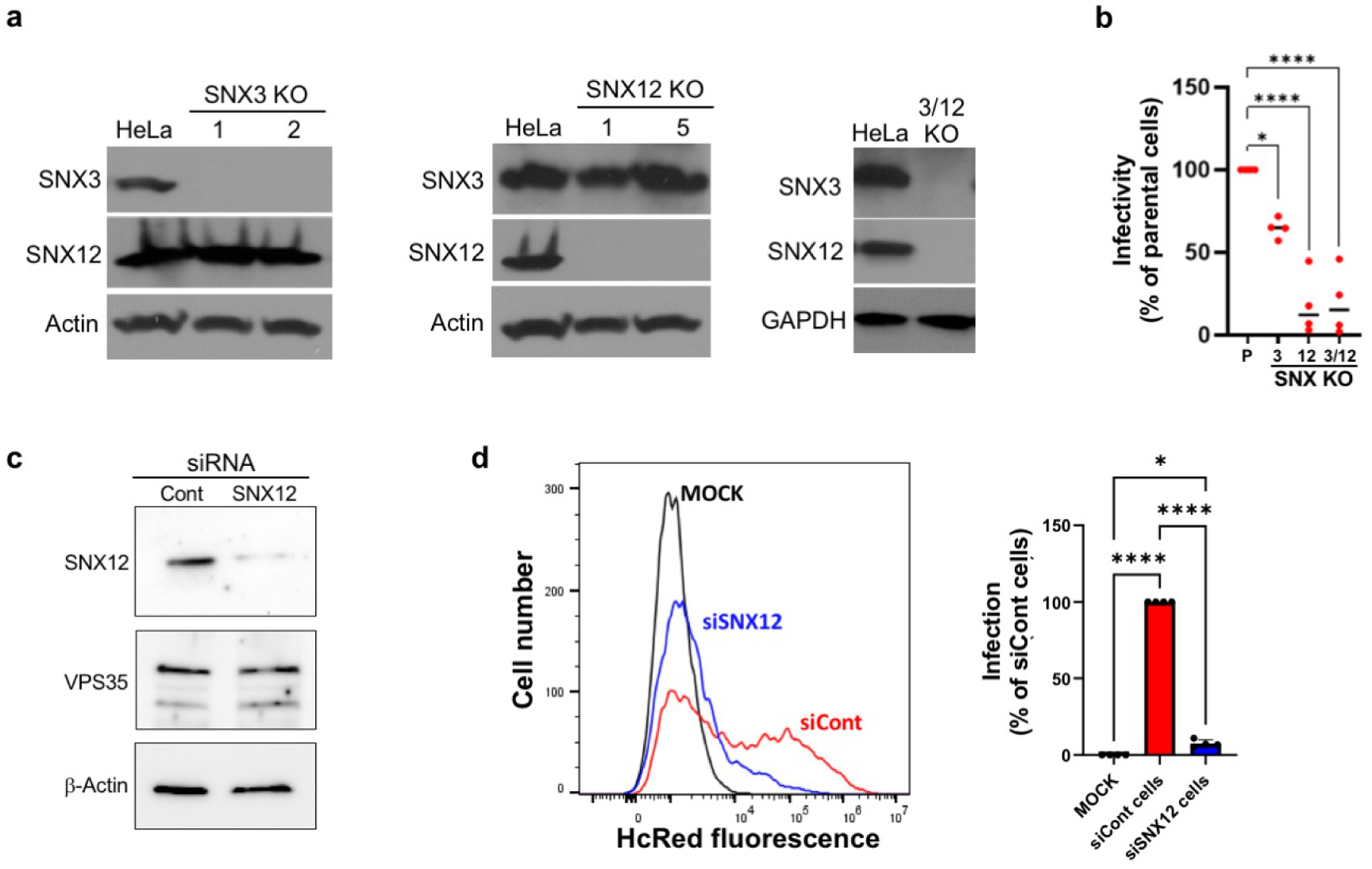
HPV requires SNX12 for efficient infection. (**a**) Independent HeLa S3 knockout (KO) cloned cell lines for SNX 3 (clone 1 and 2), SNX12 (clone 1 and 5), and for SNX3 plus SNX12 (3/12) were lysed and subjected to SDS-PAGE followed by immunoblotting with antibodies recognizing SNX3, SNX12, or -actin or GAPDH (as a loading controls). (**b**) Parental HeLa S3 cells (P) and cell clones knocked out for SNX3, SNX12, and SNX3 plus SNX12 (3/12) were infected with wild-type HPV16 PsV. At 48 hpi, cells were analyzed by flow cytometry to quantify HcRed reporter expression. The mean fluorescence intensity (MFI) of parental cells was set to 100%, and data points show the results from four independent cell lines of each genotype (as a percentage of the MFI of infected parental cells), with the means shown by the horizontal lines. Statistical significance was assessed by one-way ANOVA. Significance values: *, p<0.05; ****, p < 0.0001. (**c**) HeLa S3 cells were transfected with small interfering RNA (siRNA) targeting SNX12 or with control scrambled siRNA. 24 hours after transfection, cells were lysed and subjected to SDS-PAGE followed by immunoblotting with antibodies recognizing SNX12, VPS35, or b-actin. (**d**) SNX12 knock down cells as in panel c were mock-infected or infected with wild-type HPV16 L2-3xFLAG PsV 24 hours after transfection with control siRNA (siCont) or siRNA targeting SNX12. At 48 hpi, cells were analyzed by flow cytometry to quantify HcRed expression. The left panel shows a representative histogram, and the right panel shows the quantified infectivity for four independent experiments, expressed as a percentage of MFI of siCont infected cells, with error bars showing standard deviation. Significance values: *, p<0.05; ****, p < 0.0001

To corroborate these findings, we also used siRNAs to perform transient knockdown (KD) of SNX12. Consistent with the knockout results, siRNA-mediated depletion of SNX12 caused a marked reduction in HPV16 PsV infectivity (Fig. 1c, d). These results in KO and KD cells confirm that SNX12 is a critical and specific factor required for efficient HPV entry, whereas SNX3 appears to be less important. Based on these results, SNX12 knockdown cells were used for all subsequent cellular assays.

### SNX12 facilitates L2-retromer interactions and HPV trafficking during entry

To determine whether HPV16 and SNX12 interact during viral entry, we used the proximity ligation assay (PLA) at 8 hpi, a time when L2 has engaged retromer to exit the endosome and traffic to the TGN. This technique generates a fluorescent signal when two proteins of interest are within 40 nm. HeLa cells were infected with HPV16 PsVs containing a FLAG-tagged L2 protein and subjected to PLA at 8 hpi to assess the proximity of L2.FLAG and SNX12 (Fig. 2a). There was little PLA signal in uninfected cells, whereas infected cells treated with control siRNA exhibited a strong PLA signal. SNX12 knockdown dramatically reduced the signal as expected because it depleted one of the target proteins. These results indicate SNX12 and HPV16 L2 are in close proximity during entry at 8 hpi.

**Fig. 2:**
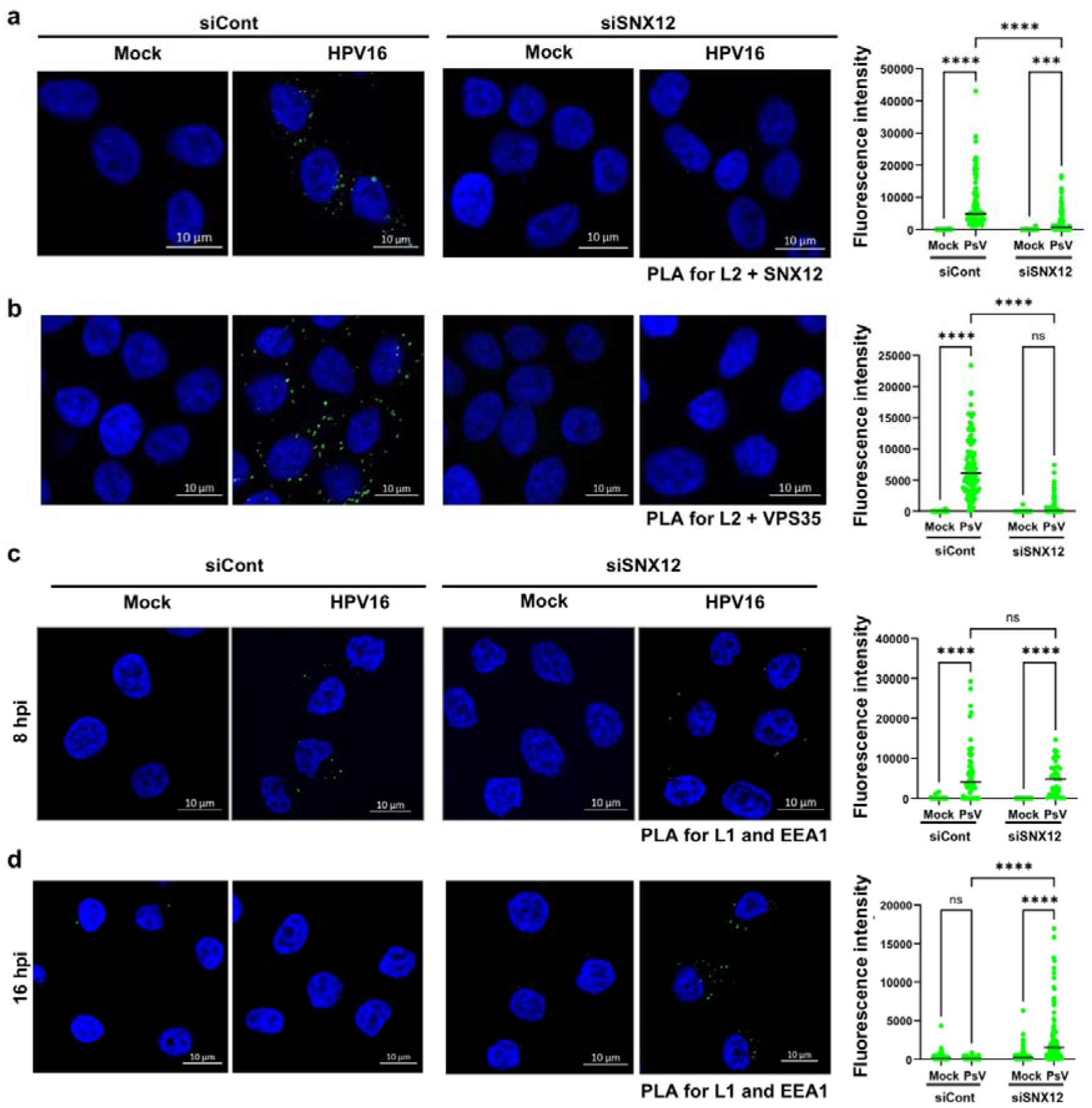
SNX12 mediates L2-retromer association and endosome escape during HPV entry. (**a**, **b**) Control- or SNX12 siRNA-treated HeLa S3 cells were mock-infected or infected with HPV16 L2-3xFLAG for 8 h and subjected to PLA assay using antibodies recognizing FLAG and either SNX12 (**a**) or VPS35 (**b**), as indicated. PLA signals are shown in green, and nuclei were stained with DAPI (blue). PLA signals in singles cells were quantified and plotted in the right panels. (**c** and **d**) Cells were mock-infected or infected as above for 8 h (**c**) or 16 h (**d**) and subjected to the PLA assay using antibodies recognizing L1 and EEA1. Results are displayed as in panel (a). In all panels, statistical significance was assessed by two-way ANOVA. Significance values: ns, not significant; ***, p < 0.001; and ****, p < 0.0001.

Upon protrusion of the C-terminus of L2 through the endosomal membrane into the cytoplasm, it binds to retromer which mediates entry of the incoming virus into the retrograde pathway. To determine whether SNX12 is required for HPV to associate with the retromer complex during infection, we performed PLA at 8 hpi with antibodies recognizing FLAG-tagged HPV16 L2 and the retromer subunit VPS35 in control and SNX12-depleted cells. As reported previously, at this time point, proximity between L2 and retromer is readily detectable in control infected cells but not in uninfected cells^23^ (Fig 2b). Notably, the L2-VPS35 PLA signal was significantly reduced in SNX12 knockdown cells, even though SNX12 KD did not affect the expression of VPS35 (Fig 1c and 2b). These results indicate that SNX12 plays an important role in recruiting retromer to incoming HPV during virus entry.

After virus internalization, HPV is first localized to the endosome, a step that does not require retromer. To determine if SNX12 KD affected arrival of HPV into the endosome during entry, we performed PLA at 8 hpi with antibodies recognizing HPV16 L1 and the early endosome marker EEA1 (Fig. 2c). There was minimal PLA signal in uninfected cells, as expected. Similar levels of L1-EEA1 PLA signals were observed in infected HeLa cells transfected with control siRNA and siRNA targeting SNX12, indicating that SNX12 depletion does not impair HPV endocytosis and that endocytosed HPV reaches the early endosome by 8 hpi. Next, we used PLA to assess HPV localization at 16 hpi. At this time during normal infection, the virus has largely exited the endosome and arrived at the TGN. HeLa cells were mock-infected or infected with HPV16.FLAG PsV, and 16 hpi PLA was performed with antibodies recognizing the L1 and EEA1. As shown in Figure 2d, there is increased L1-EEA1 signal at 16 hpi in the SNX12 KD cells compared to control cells, indicating that although HPV can enter the endosome, there is a defect in endosome exit in cells depleted of SNX12. Endosome accumulation is also observed when the retromer binding sites on L2 are removed or when retromer is knocked down^23^.

### HPV L2 C-terminus directly engages retromer-SNX12 complexes

To determine how HPV L2 engages the retromer machinery, we investigated whether the L2 C-terminal region directly interacts with specific retromer-SNX adaptor complexes. Using isothermal titration calorimetry (ITC), we measured binding affinities between a synthetic peptide corresponding to the L2 C-terminus (residues 441-460, L2(441-460)), which contains the two mapped retromer binding sites^23^, and various purified adaptor-retromer combinations, including SNX3, SNX12, ESCPE-1, SNX27, and the retromer core complex. The peptide did not bind to individual SNX proteins or retromer alone but showed direct, saturable binding to the SNX3-retromer and SNX12-retromer complexes, with dissociation constants of ∼63 μM and ∼45 μM, respectively (Fig. 3a and Extended Data Fig. 1). The higher affinity of L2 for the retromer-SNX12 complex suggests that SNX12 may provide a more stable platform for viral cargo engagement. Notably, the affinity of L2 binding to SNX12-retromer exceeded that of the endogenous cargo DMT1-II under similar assay conditions^18^, presumably reflecting the fact that viral entry pathways require high-affinity binding.

**Fig. 3:**
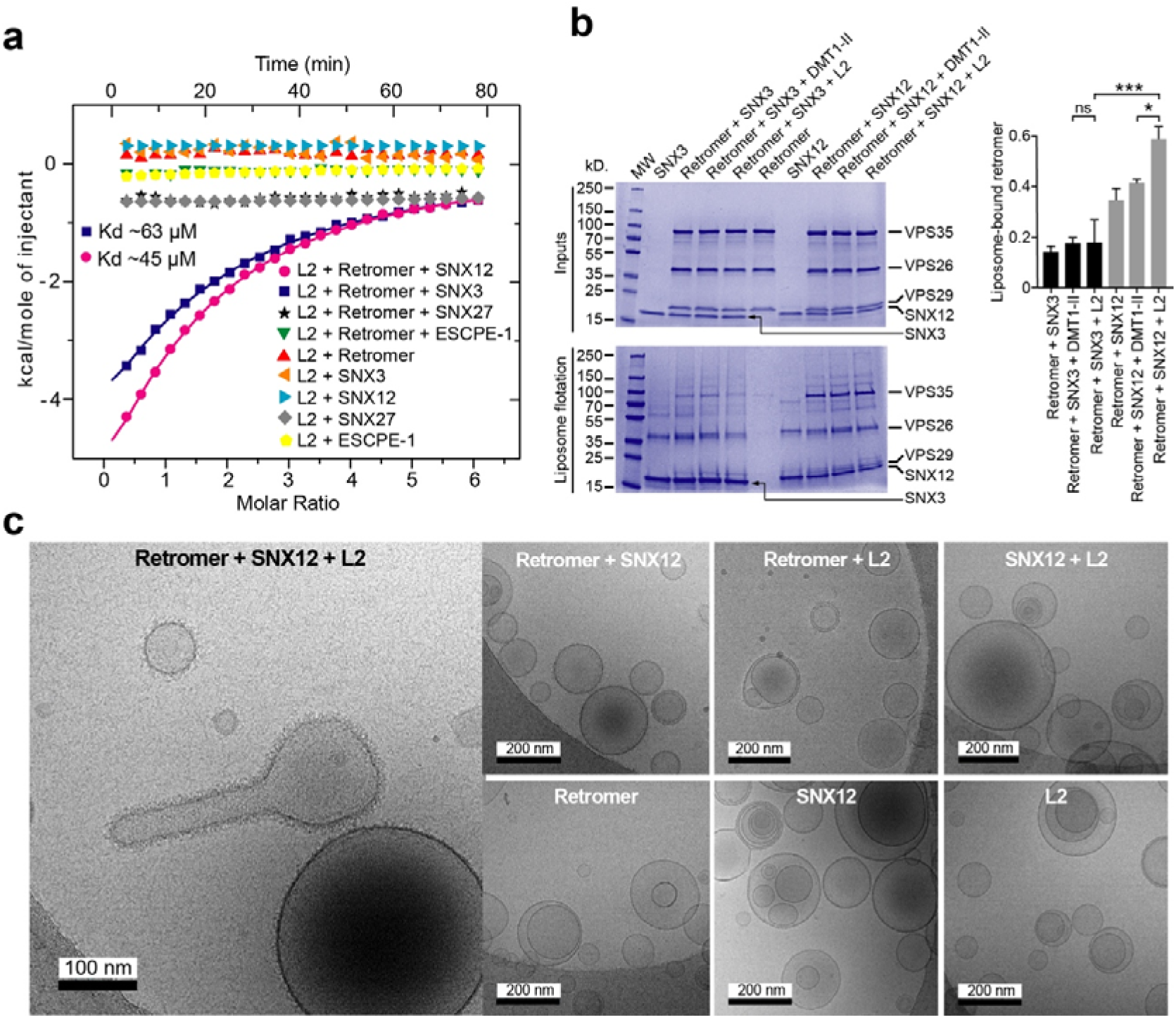
The HPV L2 tail engages retromer-PX-only SNX complexes to drive cargo-dependent membrane remodeling. (**a**) Isothermal titration calorimetry (ITC) measurements of the HPV L2(441-460) peptide binding to various retromer and retromer-SNX-associated proteins involved in different endosomal sorting routes. (**b**) Coflotation assays with retromer, SNX3, SNX12, and L2(441-460) reveal that SNX12 promotes stronger retromer association with membranes than SNX3, and is further enhanced by the HPV L2(441-460) tail. The left panel shows a Coomassie-stained gel of cofloated fractions, and the right panel quantifies retromer recruitment based on the VPS35/SNX3 or VPS35/SNX12 band intensity ratios. (**c**) Cryo-EM images showing cargo-dependent membrane tubulation. Tubule formation occurs only in the presence of retromer, an SNX adaptor, and the HPV L2(441-460) peptide.

To assess whether this interaction promotes membrane recruitment, we performed liposome flotation assays using PI3P-enriched liposomes, as SNX3 and SNX12 bound selectively to PI3P among all phosphoinositides tested (Extended Data Fig. 2a). SNX3 and SNX12 each bound PI3P-containing liposomes in the absence of retromer, consistent with PX domain-mediated recognition of PI3P, while retromer alone showed no detectable membrane association (Fig. 3b). When co-incubated with either SNX3 or SNX12, retromer was efficiently recruited to membranes; however, SNX12 promoted more robust recruitment than SNX3 (Fig. 3b). Inclusion of the L2(441-460) peptide slightly enhanced membrane association in the presence of SNX12, but not with SNX3. This modest enhancement is likely due to the unanchored and soluble nature of the peptide, which limits its ability to stably bridge retromer to the membrane. The stronger recruitment driven by SNX12 and L2 compared to SNX3 and L2 is consistent with the finding reported above that SNX12 plays a more prominent role in HPV infection.

We next asked whether L2 binding could trigger membrane remodeling. Tubulation assays using PI3P-containing liposomes revealed that SNX12 or L2(441-460) alone, or in combination with retromer, were unable to induce membrane deformation (Fig. 3c). However, the presence of retromer, SNX12 and L2(441-460) consistently triggered tubule formation (Fig. 3c). This cargo-induced remodeling was also observed with SNX3 and the endogenous retromer cargo DMT1-II (residues 550-568, DMT1-II (550-568)) (Extended Data Fig. 2b), suggesting that cargo engagement is a prerequisite for membrane tubulation by PX-adaptor:retromer complexes.

### Structural Basis of L2 Recognition by Retromer-SNX12

To determine the molecular details of how SNX12 mediates retromer recruitment to HPV, we determined the crystal structure of a ternary VPS26-VPS35N-SNX12 complex, following a strategy analogous to that used for the SNX3-retromer-DMT1-II structure^18^. The L2 C-terminal binding motif (residues 449-458) was fused to VPS26 via a flexible linker, and a synthetic peptide containing the same sequence was added during crystallization to enhance crystal quality (Fig. 4a-c). Previous studies identified two potential retromer-binding sites on HPV16 L2^23^, spanning residues ^446^FYL^448^ and ^452^YYML^455^. To assess the relative contribution of these motifs to retromer engagement, we generated two separate triple mutants: L2(441-460) F446A/Y447A/L448A targeting the FYL site and L2(441-460) Y453A/M454A/L455A targeting the YYML site. ITC analysis revealed that mutations within the YYML motif completely abolished L2 binding to the SNX12-VPS26-VPS35N complex, whereas mutations in the FYL site caused only a modest reduction in binding (Fig. 4d and Extended Data Fig. 3). Based on these results, the FYL site was not included in the L2 segment used for crystallization, as the YYML motif was clearly the dominant determinant for binding in vitro.

**Fig. 4:**
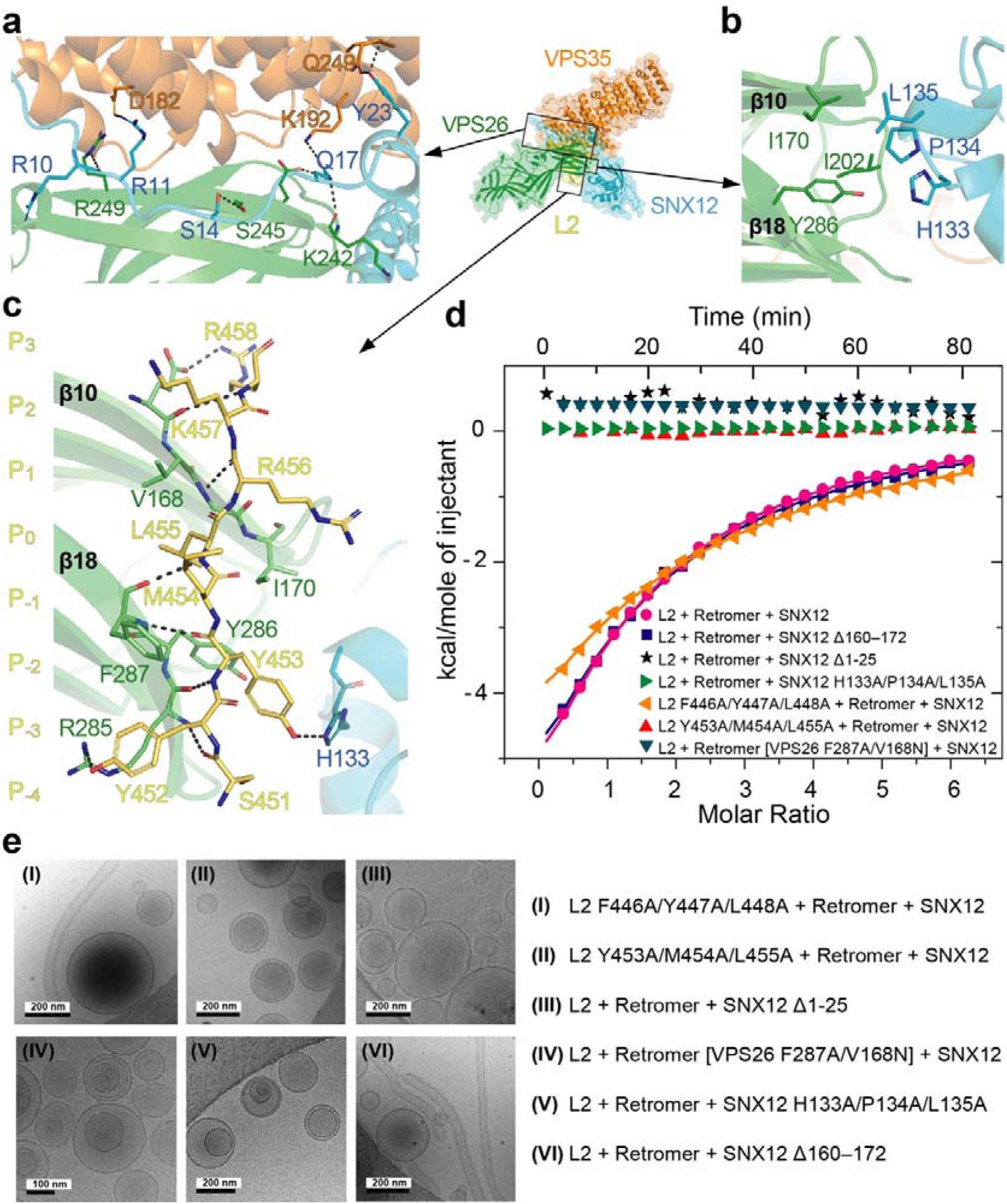
Structural basis of HPV L2 recognition by SNX12-retromer. (**a**-**c**) X-ray crystallographic model showing SNX12 contacts with the VPS26-VPS35 subcomplex and the binding of the HPV L2(449-458) at the VPS26-SNX12 interface. L2 peptide shown as stick figure, other components in ribbon diagram. (**d**) ITC analysis of HPV L2(441-460) binding to retromer-SNX12 complexes and mutants. Deletion of the SNX12 N terminus or mutations disrupting the VPS26-SNX12 interface, the VPS26 binding pocket, or the L2 YML motif (residues 453-455) abolished binding, whereas removal of the SNX12 C terminus or mutation of the FYL motif (residues 446-448) had no detectable effect. **(e)** Liposome tubulation assays showing that disruption of the SNX12 N terminus, the VPS26-SNX12 interface, the VPS26 pocket, or the L2 YML motif (453-455) suppressed tubulation, whereas mutation of the L2 FYL motif (446-448) had no appreciable effect.

The crystal structure of the retromer/SNX12/L2 complex was solved by molecular replacement and refined to 3.13 Å resolution (Fig. 4a-c, Extended Data Fig. 4a and Extended Data Table 1). The overall arrangement of SNX12, VPS26, and VPS35N closely resembled that of the previously reported SNX3:retromer:DMT1-II complex^18^, underscoring the conserved architecture of retromer assemblies formed with PX-domain adaptors (Fig. 4a-c and Extended Data Fig. 4b-d). In both cases, the PX-domain adaptor binds at the VPS26-VPS35 interface and contributes directly to the formation of the cargo-binding site on VPS26, revealing a shared structural framework for cargo recognition. The core of the L2 interaction (P_0_) is centered around L455 in the YYML binding site, which is buried within the hydrophobic pocket formed between strands β10 and β18 of VPS26. Residues M454 at P_-1_ and Y452 at P_-3_ contribute to binding by forming a hydrophobic clamp around F287 of VPS26. Additionally, Y453 at P_-2_ lies in a shallow hydrophobic cavity at the VPS26-SNX12 interface, underscoring the VPS26-SNX12 association in stabilizing the L2 interaction (Fig. 4c, and Extended Data Fig. 4b-d).

To further validate these interactions in solution, we conducted ITC and liposome remodeling assays with SNX12 and VPS26 mutants. Deletion of the first 25 residues of SNX12 (SNX12 Δ1-25), which interact with VPS26 and VPS35, or substitution of key interface residues in its PX domain (SNX12 H133A/P134A/L135A) completely abrogated L2 binding (Fig. 4d). Likewise, mutation of residues F287 and V168 in VPS26 that flank the hydrophobic binding pocket where the P_0_ residue (L455) of L2 inserts (VPS26 F287A/V168N) also abolished L2 binding (Fig. 4d). In contrast, removal of the flexible C-terminal region of SNX12 (SNX12 Δ160–172), which is not resolved in the crystal structure and is not involved in retromer assembly, had no effect on L2 binding (Fig. 4d).

Consistent with these results, liposome tubulation assays showed that the L2(441-460) Y453A/M454A/L455A mutant reduced tubulation but did not completely inhibit it (panel I in Fig. 4e), whereas mutations in the FYL site had little or no effect on tubulation (panel II in Fig. 4e). In addition, SNX12 Δ1-25, SNX12 H133A/P134A/L135A and VPS26 F287A/V168N displayed drastically reduced tubulation activity (panels III-V in Fig. 4e). In contrast, the SNX12 Δ160-172 mutant had no noticeable impact on tubulation (Panel VI in Fig. 4e). Together, these results indicate that effective membrane tubulation depends primarily on the combined contribution of the ^452^YYML^455^ motif in L2 and the structural integrity of the SNX12-retromer interface, underscoring the cooperative nature of cargo recognition and adaptor binding in coat assembly.

### Structure of the SNX12-retromer membrane coat

To elucidate the organization of SNX12-retromer assemblies on membranes, we performed cryo-electron tomography (cryoET) and subtomogram averaging (STA) on retromer-coated liposomes in the presence of L2(441-460). The initial alignment identified arch-like units on the membrane surface, which were further refined using local alignment procedures to generate an average density map at 11 Å resolution (Fig. 5a, Extended Data Fig 5 and 6, and Extended Data Table 2). At this resolution, individual subunits of the complex (VPS35, VPS26, VPS29, and SNX12) were clearly distinguishable in the STA map, allowing for the precise docking of experimental structures to assemble a pseudo-atomic model (Fig. 5a). The dimeric arch-like structure of SNX12:retromer is similar to the fungal Grd19:retromer (Grd19 is a SNX3/12 homologue) and the metazoan SNX3:retromer complex^21^, in which the VPS26 subunit and the SNX-PX domain contact the membrane at the base of each arch leg (Fig 5. a-b). Initial 3D alignment and averaging of individual sub-volumes revealed a well-defined central retromer arch, while neighboring densities exhibited partial overlapping arches, indicating mixed populations. Subsequent multi-reference alignment (MRA) revealed two distinct structural classes: class A, characterized by the presence of density corresponding to a neighbouring retromer arch (“arch”), and class V, which at the equivalent position lacks arch density and instead represents the valley between two arches (“valley”) (Fig. 5c-e and Extended Data Fig. 7). In class A, the neighboring plane intersects an adjacent arch, whereas in class V, the neighboring plane intersects the region between two arches (Extended Data Fig 7). By contrast, in the other lateral plane both classes display an identical arch arrangement relative to the central arch reference (Fig. 5e), suggesting that retromer may assemble in a patterned manner along the tubule.

**Fig. 5:**
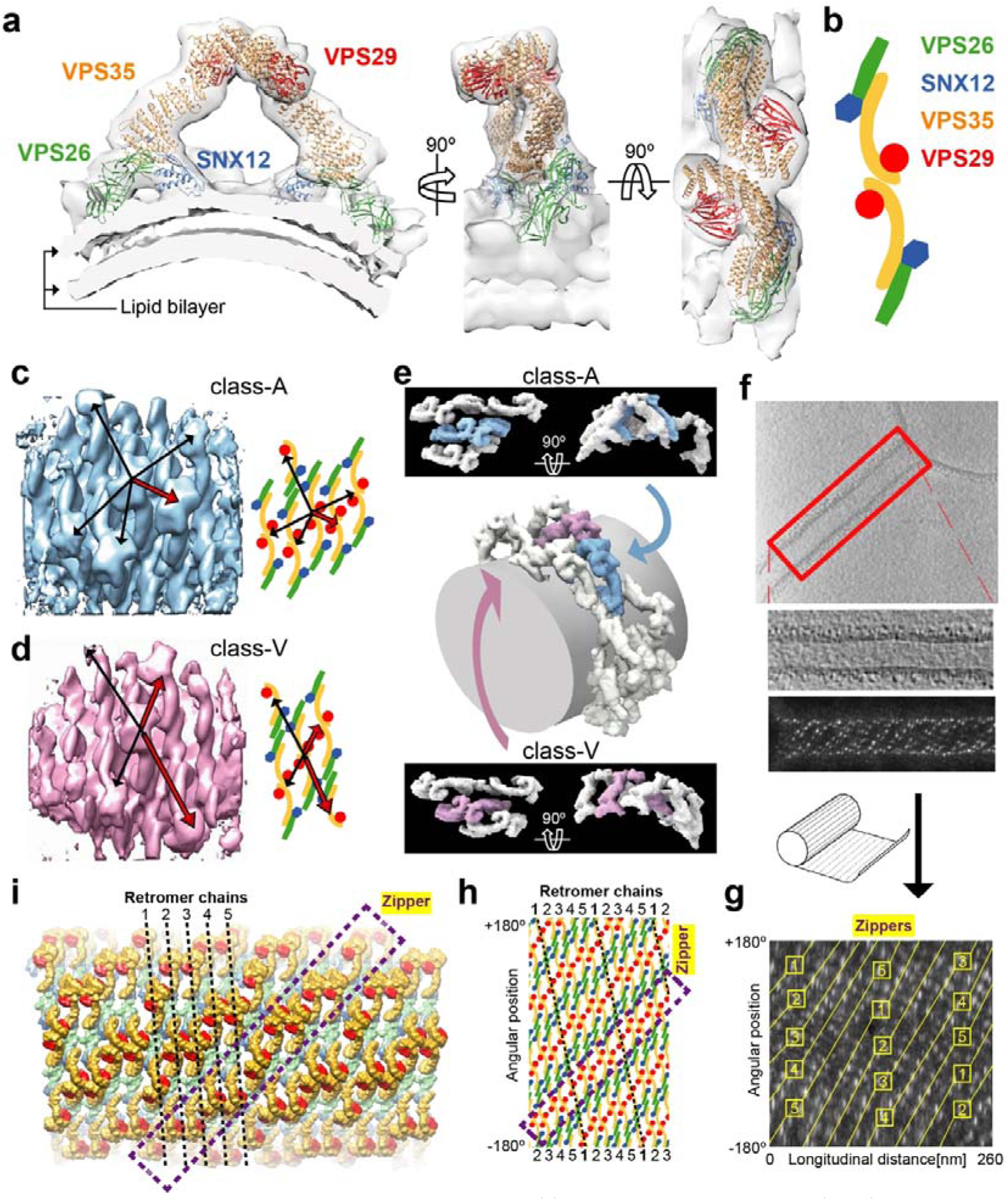
Architecture of the retromer-SNX12 coat. (**a**) Subtomogram-averaged (STA) structure of the retromer-SNX12 arch-like dimer, showing docked components from the X-ray models (PDB 9Q8M, this work, and PDB 2R17^50^. (**b**) Schematic representation of the retromer-SNX12 arch components viewed from the top. (**c**) Structure and schematic representation of the class-A neighborhood. Arrows indicate the nearest retromer arches relative to the central arch. Red arrow points to the neighbouring retromer arch. (**d**) Corresponding view of the class-V neighborhood were red arrows point to the valley between two arches (**e**) Specific arch positions identified by template matching, illustrating the coexistence of class-A and class-V conformations in the 5-start model. (**f**) Representative cryo-electron tomogram showing a retromer-coated membrane tubule emanating from a liposome (top). A magnified view of the tubule highlights retromer assemblies decorating the membrane surface (middle). The bottom panel shows a template-based maximum intensity projection of the assembled cross-correlation map, viewed from the side of the tubule, revealing the pseudo-periodic organization of retromer arches along the membrane. (**g**) “Unrolled” view of panel (**f**), highlighting the “zipper-like” arrangement of paired arches. (**h**, **i**) 2D (**h**), and 3D (**i**) representation of an idealized 5-start helical model of the retromer-SNX12 coat from (**g**) defined by a 10° chain angle, a tube radius of 27 nm, and an arc-to-arc distance of 26 nm.

**Fig. 6:**
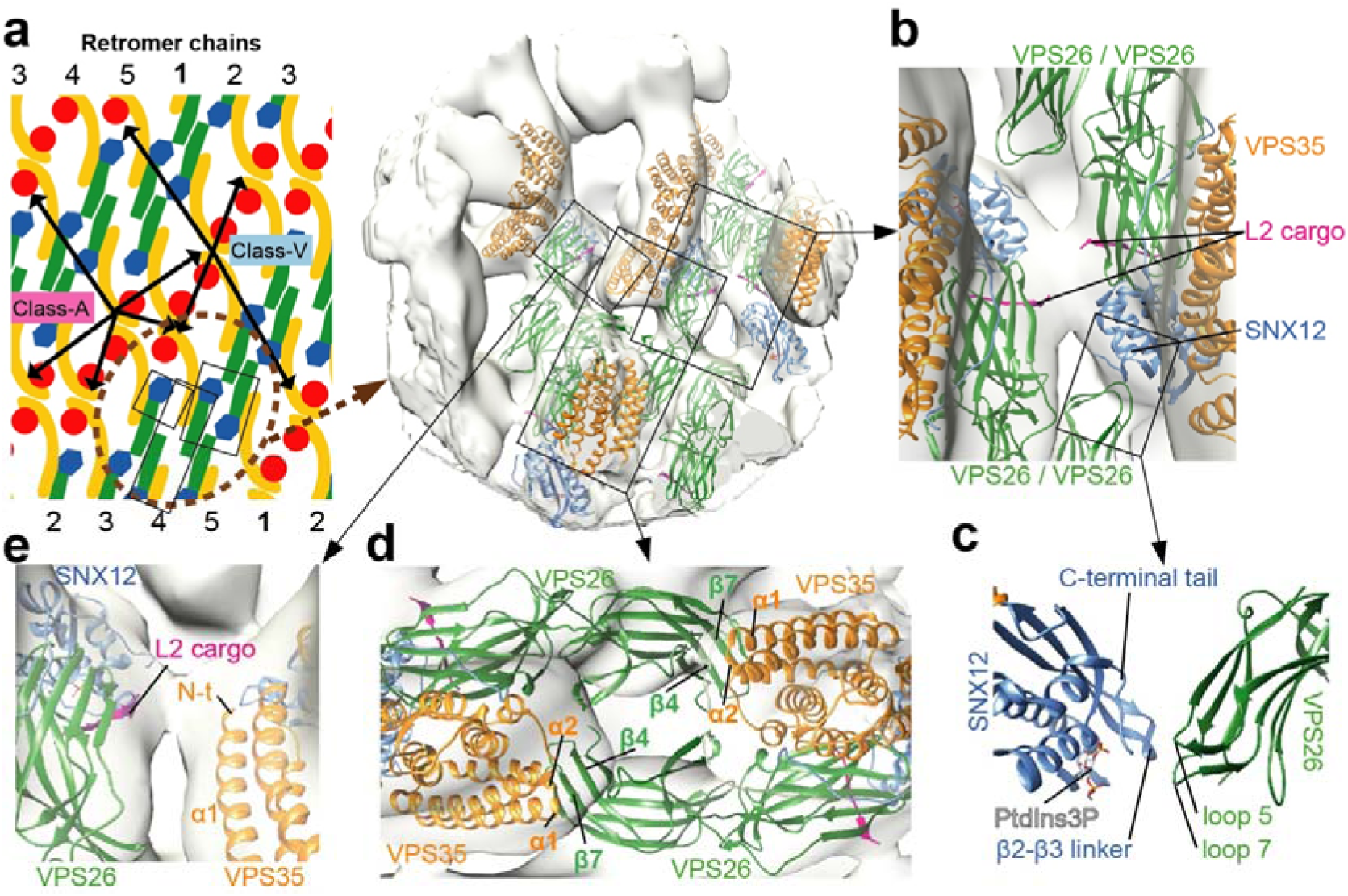
Lateral interaction regions in the retromer-SNX12 coat. (**a**) Close-up view of the 5-start helical coat highlighting the organization of neighboring class-A and class-V arches. (**b**, **c**) Interaction between the SNX12 PX domain and the N-terminal lobe of VPS26. Note in (B) the close apposition of two L2-binding interfaces bridged by continuous electron density. (**d**) Intertwined association between two VPS35 subunits docked onto the N-terminal lobes of adjacent VPS26 molecules. (**e**) VPS26 homodimerization within laterally associated retromer units positions the L2-binding site adjacent to the N-terminal tail of a neighboring VPS35, with continuous density bridging the interface.

Next, to construct a detailed landscape of the retromer-SNX12 coat, we performed template matching of retromer arches on several representative tomograms. Despite distortions in the tubular coat organization and the absence of long-range axial symmetry, a pseudo-periodic pattern emerged (Fig. 5f-i). This pattern corresponded to right-handed helical chains (retromer chains), defined as retromer arches periodically connected at the base through the dimerization of the N-terminal domains of Vps26. These chains wrapped around the membrane, with either five or six helical starts depending on the tube (Fig. 5 g-i, Extended Data Fig. 8, Extended Data Movie 1 and Extended Data Movie 2). The lateral packing of retromer arches along the longitudinal axis of the tube exhibited a left-handed ‘zipper’ organization in both the 5-start and 6-start helices, with distinct patterns of class-A and class-V. In the modelled 6-start helix, retromer arches exhibited only a class-A distribution, whereas in the 5-start helix model, they displayed an alternating arrangement of class-A and class-V (Fig. 5h-i, Fig 6a, and Extended Data Fig. 8). Together, these analyses revealed that in the presence of the L2 peptide retromer-SNX12 forms a conserved arch structure that organizes into pseudo-periodic helical coats along membrane tubules, exhibiting distinct spatial patterns.

### Lateral contacts in the membrane-proximal region contribute to coat organization

In both class-A and class-V subtomograms, the legs and apex of the retromer arches showed no lateral contacts, suggesting that coat organization is primarily governed at the membrane-proximal level. To investigate this further, we performed focused alignment and averaging on the base of the central arch in class A of the 5-start helix, as this region includes two side-facing areas that encompass all lateral contacts observed in both classes (Fig. 6a). Although the resolution was insufficient to pinpoint specific interacting residues, the presence of continuous electron density bridging adjacent subunits at distances below 10–15 Å enabled the characterization of contact regions. One such contact involves loops 5 and 7 of VPS26, along with the N-terminus of VPS26, which interact with the PX domain of SNX12 at the β2-to-β3 linker and C-terminal tail, adjacent to the PtdIns3P-binding pocket (Fig. 6b, c). Through this interaction, an assembly of alternating VPS26 and SNX12 subunits emerges, with the N-terminal and C-terminal lobes of VPS26 engaging opposite faces of the SNX12 PX domain. Remarkably, this alternating organization and VPS26 homodimerization create a geometry that positions two L2 cargo-binding interfaces in close apposition, with continuous electron density bridging them, suggesting that L2 may strengthen the lateral interactions between adjacent retromer arches. (Fig. 6b).

Another contact involves an intertwined association between two retromer complexes, in which helices α1 and α2 of the VPS35 N-terminus dock onto strands β4 and β7 on the convex surface of the VPS26 N-terminal lobe (Fig. 6d). Interestingly, an interaction bearing a similar pattern has been previously resolved in the absence of membranes by single-particle cryo-EM^26^. In this configuration, homodimerization of VPS26 with a laterally associated retromer positions the L2 cargo-binding site adjacent to the N-terminal tail of a nearby VPS35, with continuous electron density bridging the interface (Fig. 6e). These observations indicate that lateral retromer interactions are confined to the membrane-proximal layer, where contacts between VPS26, SNX12, and VPS35 bring cargo-binding sites into close proximity. These findings suggest that cargo may play a dual role, both promoting retromer-SNX12 assembly and bridging adjacent chains to stabilize the coat.

### Hinges at VPS26 sites and multi-start helices underlie coat adaptation to variable membrane diameters

Our liposome-tubulation assays produced tubes with diameters ranging from approximately 40 to 60 nm, measured across the lipid bilayer (Fig. 7a). To examine how the SNX12–retromer coat accommodates different membrane diameters, we focused on the two predominant tubule populations, with radii of ∼22 nm and ∼26 nm, which together accounted for more than 50 % of all observed tubules (Fig. 7a, b). Smaller-diameter tubules displayed 5 helix starts with a helical pitch of ∼11.4°, whereas larger-diameter tubules exhibited 6 helix starts and a slightly reduced helical pitch of ∼10.0°. The Euclidean distance between adjacent VPS26 dimers remained relatively constant (∼27.2 nm in small tubes vs. ∼27.9 nm in large tubes), indicating that local spacing between arches is maintained. To further explore structural adaptations, we computed angles and distances using equivalent reference points for both tubule types. These included the VPS26-VPS26 contacts between arches, the VPS35-VPS35 contact at the apex of the arch, and a point beneath the center of the arch where it intersects the membrane. This comparative analysis revealed that at the VPS26-VPS26 inter-arch contacts, the angle between adjacent arches decreased by ∼8° in larger-diameter tubules (Fig. 7c), while the apex angle at the VPS35-VPS35 interface remained constant (∼85°) (Fig. 7d and Extended Data Movie 3). This angular expansion is consistent with a flexible hinge mechanism at the base of the coat, allowing adjacent arches to tilt apart while preserving the structural integrity of each individual arch. These findings support the notion that retromer adapts to membrane curvature via helical variation and hinge flexibility at the membrane-proximal region while preserving the arch geometry.

**Fig. 7:**
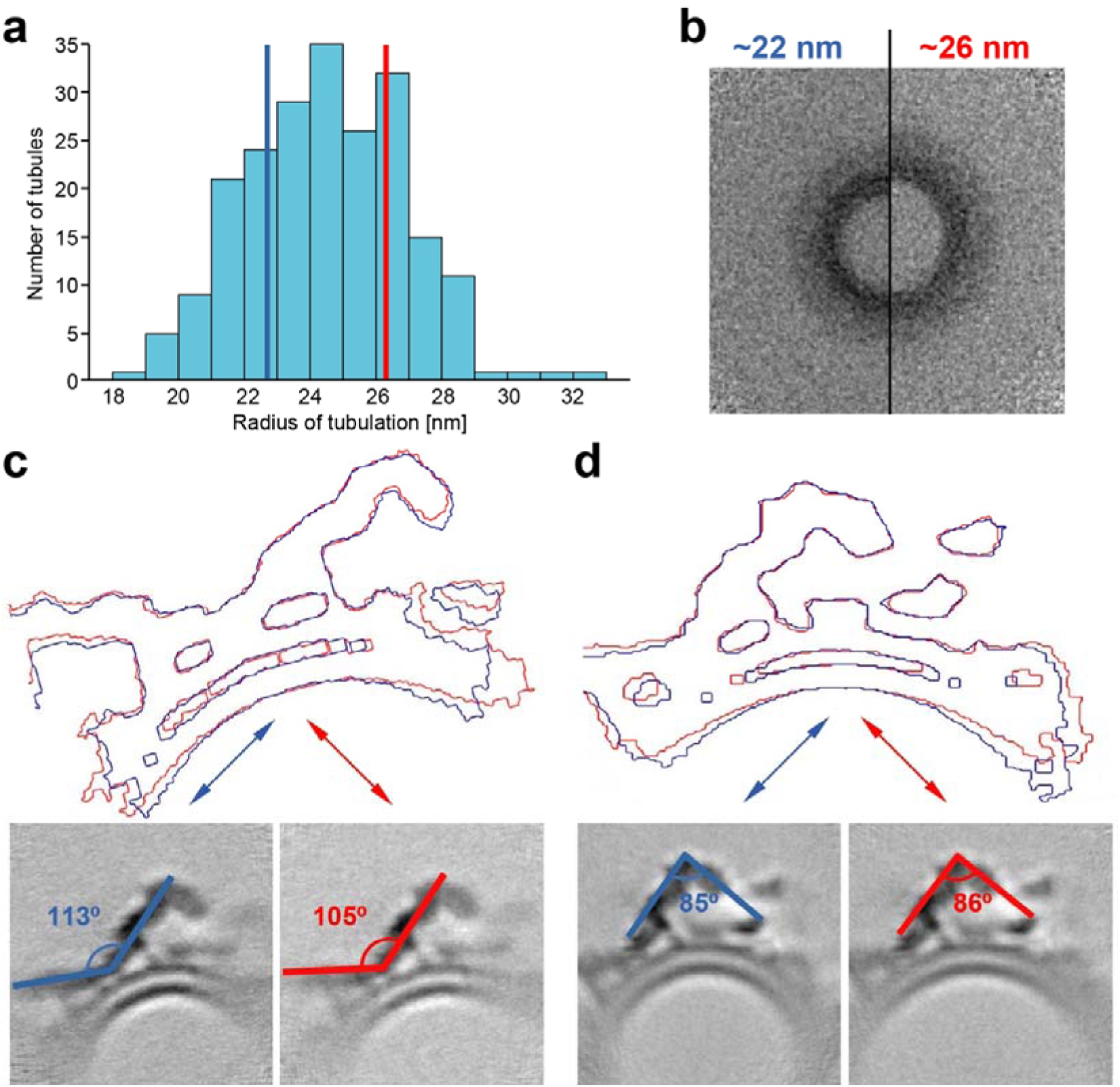
Hinge flexibility at inter-arch contacts enables coat adaptation to membrane curvature. (**a**) Distribution of tubulation radii, with thresholds defining small (blue) and large (red) tubule populations. (**b**) Half cross-sections of averaged small (right) and large (left) tubule populations showing differences in diameter. (**c**) Contour and angular comparison of retromer-SNX12 assemblies in a slice containing a VPS26-VPS26 inter-arch contact. (**d**) Equivalent analysis for a slice containing a VPS35-VPS35 contact representing the dimeric retromer arch.

## Discussion

The results presented here resolve the identity of the SNX adaptor that cooperates with retromer to drive HPV16 endosomal trafficking, and in doing so reveal a new principle of retromer coat architecture; the combination of SNX adaptor and cargo co-determines the higher-order geometry of the coat lattice.

SNX12 binds via retromer near the C-terminus of L2^22,23^, a position likely to become accessible earlier during translocation than the more N-terminal motifs recognized by SNX17 and SNX27^27,28^, thus facilitating timely engagement of the retrograde pathway toward the TGN. Given the high sequence homology between SNX12 and the more abundant SNX3^29^, the selective use of SNX12 by HPV16 was unexpected. At the level of cargo recognition, the selectivity is not immediately obvious: L2 binds the conserved hydrophobic groove of VPS26 while contacting the SNX PX domain in a mode closely related to that used by the endogenous cargo DMT1-II^18^, and L2 exhibits only slightly higher affinity for SNX12-retromer than for SNX3-retromer. Consistently, in vitro reconstitution shows that both L2 and DMT1-II can trigger membrane tubulation with either SNX adaptor, demonstrating that short retromer-binding motifs are sufficient to drive coat assembly and membrane remodeling regardless of adaptor identity.

The structural rationale for SNX12 selectivity instead emerges from cryo-electron tomography. Retromer assembles into arch-shaped dimers that form helical lattices around membrane tubules, but the geometry of this lattice is adaptor dependent. Whereas SNX3–retromer adopts a single pseudohelical arrangement^21^, SNX12-retromer forms two distinct arch-packing environments, class A and class V, that organize into patterned five- or six-start helices, revealing a distinct coat architecture. In fact, this dual organization is not a consequence of sample heterogeneity but rather an intrinsic property of the coat, encoded by specific lateral contacts formed at the SNX12-VPS26 and VPS35-VPS26 interfaces. Our structural analysis is consistent with L2 peptides cross-linking adjacent arches, suggesting that cargo acts as an integral structural component of the coat rather than merely a recruitment signal. These observations indicate that adaptor-cargo identity can directly influence coat geometry and suggest that the architecture established by SNX12-retromer is particularly suited to support HPV entry. Notably, dimerization of the retromer cargo receptor SORLA has been shown to reinforce retromer coat stability^30^, raising the possibility that SORLA may similarly cross-link adjacent arches to stabilize the lattice during endosomal sorting.

These findings reframe retromer not as a fixed scaffold but as a compositionally tunable system capable of generating architecturally distinct carriers for different sorting events. Adaptor and cargo identity may therefore act as geometric inputs to coat assembly, providing a general mechanism by which cells diversify membrane traffic without expanding the repertoire of core coat proteins, a principle exploited by pathogens such as HPV through the SNX12 pathway.

## Methods

### Peptides

All synthetic peptides were purchased from Genscript (purity ≥95%) and dissolved in the buffer appropriate for each experiment. The following peptides were used: L2(441-460) (ADAGDFYLHPSYYMLRKRRK), L2(449-458) (HPSYYMLRKR), L2(441-460) F446A/Y447A/L448A (ADAGDAAAHPSYYMLRKRRK), L2(441-460) Y453A/M454A/L455A (ADAGDFYLHPSYAAARKRRK), and DMT1-II(550-568) (AQPELYLLNTMDADSLVSR).

### Lipids

All synthetic lipids were purchased from Avanti Polar Lipids in powdered format. Phospholipids (DOPC, Cat# 850375P; DOPE, Cat# 850725P; DOPS, Cat# 840035P; rhodamine-PE, Cat# 810150P) were dissolved in chloroform. Phosphoinositides [PI(3)P, Cat# 850150P; PI(4)P, Cat# 850151P; PI(5)P, Cat# 850152P; PI(3,4)P2, Cat# 850153P; PI(3,5)P2, Cat# 850154P; PI(4,5)P2, Cat# 850155P; PI(3,4,5)P3, Cat# 850156P] were resuspended in a CHCl₃:MeOH:H₂O mixture at a 20:9:1 molar ratio.

### Cell lines

HeLa S3 cells were obtained from the American Type Culture Collection (ATCC), and HEK 293TT cells were gift from Christopher Buck (NIH). All cell lines were maintained at 37°C with 5% CO₂ in Dulbecco’s Modified Eagle’s Medium (DMEM) supplemented with 20 mM HEPES, 10% fetal bovine serum (FBS), L-glutamine, and 100 units/mL penicillin-streptomycin (DMEM10). Where possible, cells were verified by the ATCC cell line authentication service.

### Recombinant DNA Procedures

All the cDNAs used in this study encode for human proteins and were cloned into bacterial expression vectors. DNA sequences encoding full-length SNX3 and SNX12, the two SNX12 deletion mutants, SNX12(Δ160-172) and SNX12(Δ1-25) and the mutant SNX12 H133A/P134A/L135A were cloned into pHis-MBP-Parallel2 vector ^31^ to express these proteins with a cleavable N-terminal 6xHis-maltose binding protein (MBP) tag. For generation of point mutations, SNX12 was cloned by Gibson assembly ^32^ into the same vector. DNA encoding SNX1, SNX27, VPS26A and the fusion construct of VPS26A with the C-terminal tail of L2, VPS26A-L2(449-458) were cloned using pET28-Sumo3 vector (EMBL, Heidelberg) with a cleavable 6xHis-Sumo3 tag at the N terminus. The N-terminal segment of human VPS35 (residues 14-470), full-length VPS35, VPS29 and SNX5 were cloned into pGST-Parallel2 vector ^31^ with a cleavable N-terminal Glutathione S-transferase (GST) tag. All constructs were verified by DNA sequencing.

### Establishment of SNX3 KO, SNX12 KO, and SNX3/12 KO cells

Guide RNAs (gRNAs) targeting SNX3 or SNX12 were designed by using the CRISPR design tool (http://crispr.mit.edu/) ^33^. Oligonucleotides encoding gRNAs targeting the human SNX12 gene (sense: CACCGACTGTAGCGCCGCCGTACGC; antisense: AAACGCGTACGGCGGCGCTACAGTC) and the human SNX3 gene (sense: CACCGGGGTCCGTAGGCGTCATTC; antisense: AACGAATGACGCCTACGGACCCC, ^18^ were purchased from IDT, annealed, and cloned into the lentiCRISPR v1 plasmid (Addgene #49535) to generate lentiCRISPR-sgSNX3 and lentiCRISPR-sgSNX12, respectively.

Lentiviruses were produced by seeding 293T cells in 12-well plates and co-transfecting them with 0.8 µg of lentiCRISPR-sgSNX3 or lentiCRISPR-sgSNX12, 0.4 µg of the lentiviral packaging plasmid psPAX2, and 0.4 µg of the plasmid pMD2.G expressing the vesicular stomatitis virus G protein. Forty-eight hours post-transfection, the lentiviral supernatant was collected, filtered, and stored at -80°C.

Knockout cells were generated by infecting HeLa cells with lentivirus and selecting them for 48 hours in medium containing 1 µg/mL puromycin. After selection with puromycin, single cells were plated in 96-well plates, and monoclonal cell lines were isolated and validated by Western blotting. Cells doubly knocked out for SNX3 and SNX12 were created by infecting SNX3 KO cell lines with lentiviruses containing the SNX12gRNA sequence. The successful double KO monoclonal cell lines were validated by Western blotting. SNX12 knock down cells were generated by using lipofectamine RNAimax to transfect 10 nM siRNAs targeting SNX12 (siGENOME Human SNX12 siRNA, 29934, Dharmacon) into Hela S3 cells. Antibodies used for western blotting were anti-SNX12 (Santa Cruz sc-101308), anti-SNX3 (Abcam ab56078), VPS35 (Abcam ab157220), anti- β-actin (Sigma, A5441-2ML), anti-pan-actin (Cell Signaling Technologies 4968S), and anti-GAPDH (sc-47724).

### Production of HPV pseudovirus (PsV)

HPV16 PsVs containing three tandem copies of the FLAG epitope tag at the C terminus of L2 were generated by co-transfecting 293TT cells with p16SheLL-3XFLAG plasmid ^34^ and reporter plasmid pCAG-HcRed (Addgene #11152) using polyethyleneimine (MilliporeSigma). Seventy-two hours post-transfection, the cells were harvested and incubated overnight at 37°C with lysis buffer (DPBS containing 0.5% Triton X-100, 10 mM MgCl₂, 5 mM CaCl₂, and 100 μg/mL RNAse A (Qiagen)) to allow capsid maturation. The clarified lysates containing matured PsVs were loaded onto an OptiPrep gradient and centrifuged at 50,000 × g for 4 hours at 4°C using a SW-55Ti rotor (Beckman). Fractions were collected and analyzed by SDS-PAGE followed by Coomassie blue to assess the abundance of L1. Peak fractions banding at the density of full capsids were pooled, aliquoted, and stored at -80°C.

### Infectivity

1 × 10⁵ control or KO HeLa S3 cells seeded in 12-well plates were incubated with wild-type HPV16 PsVs at an infectious multiplicity of infection (MOI) of approximately 2. The infectivity was determined at 48 hpi by flow cytometry analysis of HcRed fluorescence following PsVs infection. For siRNA experiments, 5 × 10⁴ HeLa S3 cells were seeded in 12-well plates and transfected with 10 nM siRNA targeting SNX12. After 24 h, cells were infected with HPV16.3×FLAG PsV at an MOI of approximately 2. Infectivity was determined as above.

### Proximity ligation assay

2.5 × 10^4^ HeLa S3 cells seeded on glass coverslips in 24-well plates were transfected with 10 nM siRNA for 24 h and then infected with HPV16-3X-FLAG PsV at MOI of 200. After 8 or 16 hours, cells were fixed with 4% paraformaldehyde and permeabilized with 0.1% saponin for 15 minutes at room temperature. Cells were then incubated with pairs of mouse and rabbit antibodies at 4°C overnight. PLA was carried out with Duolink reagents from Sigma Aldrich according to the manufacturer’s directions as described ^35^. Briefly, cells were incubated with a pair of suitable PLA antibody probes in a humidified chamber, and then subjected to ligation and amplification with fluorescent substrate at 37°C. Antibodies used were anti-HPV16 L1 (BD Biosciences 554171), anti-FLAG (for FLAG-tagged L2, Sigma F1804-1MG), anti-SNX12 (ThermoFisher PA5-99046), anti-VPS35 (Abcam ab157220), and anti-EEA1 (Cell Signaling Technology 2411S). Slides were mounted in mounting solution with DAPI. Cells were imaged with Zeiss 880 or 980 confocal microscope. Images were processed and quantified by Fiji to measure fluorescence intensity in each sample.

### Recombinant protein expression

The following proteins were expressed in *Escherichia coli* BL21 (DE3) strain: VPS35, VPS29, VPS26A, VPS26A-L2(449-458), VPS35(14-470), SNX1, SNX3, SNX5, SNX12 and SNX27. Full-length SNX12, SNX12(Δ160-172), SNX12(Δ1-25) and SNX12 H133A/P134A/L135A were expressed in *E. coli* Rosetta (DE3) strain. Both strains were grown in Luria-Bertani (LB) broth at 37°C, and protein expression was induced with 0.5 mM isopropyl-β-_D_-thiogalactopyranoside (IPTG) when the optical density at 600 nm (OD_600_) reached 0.6-0.8. Cells were harvested after 16-20 hr of growth at 18°C.

### Protein purification

All purification steps were performed at 4°C. Full-length retromer (VPS26-VPS35-VPS29), VPS35(14-470)-VPS26A-L2(449-458) complex and SNX3 were isolated following the same protocol as in Lucas et al., ^18^, whereas SNX1-SNX5 complex was purified as in López-Robles et al., 2023 ^36^.

For the purification of SNX12 WT, SNX12(Δ160-172), SNX12(Δ1-25) and SNX12 H133A/P134A/L135A, the cell pellet was lysed by sonication in buffer A [300 mM NaCl, 1 mM dithiothreitol (DTT), 20 mM imidazole, 50 mM Tris-HCl pH 8.0] supplemented with 0.5 mM phenylmethylsulfonyl fluoride (PMSF), 5 mM benzamidine, DNAses and lysozyme. After centrifugation at 5,000 xg for 45 min, the soluble fraction was incubated for 2 h in batch with Ni^2+^-nitrilotriacetate (NTA) agarose resin (QIAGEN). After extensive washing of the beads with buffer A in a gravity column, protein was eluted in buffer B (150 mM NaCl, 1 mM DTT and 250 mM imidazole, 50 mM Tris-HCl pH 7.5). Tobacco etch virus (TEV) protease was added to the eluted sample and was incubated for 3 hr to remove the N-terminal 6xHis-MBP tag. Mixture of protein and protease was dialyzed overnight in buffer C (120 mM NaCl, 1 mM DTT, 25 mM Tris-HCl pH 6.5). Proteins were finally purified by ion-exchange chromatography (HiTrap SP, GE Healthcare) using a linear NaCl gradient from 100 mM to 1 M over 20 column volumes, followed by size-exclusion chromatography (Superdex 75 16/60, GE Healthcare) in buffer D (300 mM NaCl, 1 mM DTT, 25 mM Tris-HCl, pH 7.5).

Initial steps for SNX27 purification were carried out as for SNX12, but the protein was eluted in buffer A supplemented with 250 mM imidazole and Sentrin-specific protease 2 (SENP2) was added to the eluted protein for the cleavage of the 6xHis-Sumo3-tag during an overnight dialysis in buffer E (150 mM NaCl, 1 mM DTT, 25 mM Tris-HCl pH 8.0). SNX27 was then purified by ion-exchange chromatography (HiTrap Q, GE Healthcare) using a linear NaCl gradient from 50 mM to 500 mM over 20 column volumes, followed by size-exclusion chromatography (Superdex 200 16/60, GE Healthcare) in buffer D. The concentration of all purified proteins was calculated using the theoretical extinction coefficient obtained from the Expasy ProtParam tool (https://web.expasy.org/protparam/).

### Isothermal Titration Calorimetry Assays

Isothermal titration calorimetry (ITC) experiments were conducted at 25°C on a MicroCal PEAQ-ITC instrument (Malvern Panalytical). Proteins and synthetic peptides were dialyzed overnight at 4°C against 300 mM NaCl, 0.5 mM TCEP-NaOH, 50 mM HEPES 7.5. Dialyzed proteins were degassed for 10 min in a ThermoVac sample degasser before titration. Each experiment involved one initial 0.5 μl injection (discarded in data fitting) followed by 24 injections of 1 μl aliquots with a spacing of 180 s between them. Similar injections of peptides in buffer were carried out to determine the heat of dilution of each one and the background was subtracted from the experimental data. Data were integrated and fitted to a one-site model using the Origin ITC 7.0 software package. Kinetic values as binding constant (K_a_, K_d_ =1/K_a_), stoichiometry (n) and binding enthalpy (ΔH) were obtained directly from the fit. For the analysis of the interaction between L2(441-460) and retromer in presence of SNX3 or SNX12, 960 μM of a cargo L2(441-460), L2(441-460) mutants, DMT1-II(550-568), DMT1-II Y555A/L557A or CI-MPR(2347-2375) was titrated into the calorimetric cell containing 20 μM retromer complex, or 200 μM SNX (SNX3, SNX12, SNX27 or SNX1-SNX5), or 20 μM retromer plus 200 μM SNX (SNX3, SNX12, SNX12(1-159), SNX12(26-172), SNX12 H133A/P134A/L135A or SNX27. For each condition, at least three independent ITC experiments were performed.

### Liposome Preparation

Liposomes for flotation assays were prepared by mixing DOPC, DOPE, DOPS, a phosphoinositide, and rhodamine-PE at a molar ratio of 45:28:20:5:2, respectively. For negative controls lacking phosphoinositides, liposomes were prepared with DOPC, DOPE, DOPS, and rhodamine-PE at a ratio of 47:29:22:2. Lipid mixtures were incubated at 37°C for 1 h to ensure homogenization, dried under vacuum, and rehydrated in flotation buffer A (FB-A: 150 mM NaCl, 0.5 mM TCEP, 10 mM HEPES pH 7.5, 10% sucrose) to a final lipid concentration of 1 mM. The resuspended mixtures were heated at 60°C for 1 h, subjected to 11 cycles of freeze-thaw-vortexing (liquid nitrogen and 50°C water bath), and extruded 11 times through a 0.2 μm polycarbonate filter using a mini-extruder (Avanti Polar Lipids) to obtain unilamellar vesicles. Liposomes for tubulation assays were prepared similarly by mixing DOPC, DOPE, DOPS, and PI(3)P at a 45:30:20:5 molar ratio, at a final concentration of 1.5 mM in tubulation buffer (150 mM NaCl, 1 mM TCEP, 20 mM HEPES pH 7.5). Extrusion was carried out through a 0.2 μm polycarbonate filter (Avanti Polar Lipids). Liposomes were stored at 4 °C under argon and used within three days.

### Liposome Flotation

A total of 100 μL of protein solution in flotation buffer B (FB-B: 150 mM NaCl, 0.5 mM TCEP, 10 mM HEPES pH 7.5) containing either 25 μM SNX (SNX3, SNX12, SNX12(1-159), SNX12(26-172), or SNX12 H133A/P134A/L135A), 25 μM retromer, 25 μM SNX plus 25 μM retromer, or 25 μM SNX plus 25 μM retromer supplemented with 187.5 μM L2(441-460) or DMT1-II(550-568) peptide, was mixed with 100 μL of 1 mM liposomes prepared in FB-A and incubated for 30 min at 4°C. Each mixture was combined with 100 μL of flotation buffer C (FB-C: 150 mM NaCl, 0.5 mM TCEP, 10 mM HEPES pH 7.5, 80% sucrose) to yield a final 30% sucrose layer, which was placed at the bottom of an Ultra-Clear™ centrifuge tube (Beckman Coulter, ref. 344090). A discontinuous sucrose gradient was formed by sequentially overlaying 300 μL of flotation buffer D (FB-D: 150 mM NaCl, 0.5 mM TCEP, 10 mM HEPES pH 7.5, 25% sucrose) and 50 μL of FB-B. Samples were centrifuged at 44,000 rpm for 1 h at 4°C in a SW55Ti rotor (Beckman Coulter). Following centrifugation, the liposome-containing (pink-rhodamine) fraction at the interface between 25% and 0% sucrose was collected. Liposome recovery was quantified by measuring rhodamine absorbance at 574 nm, and samples were normalized to the condition with the lowest liposome content. Proteins were analyzed by SDS-PAGE on 4–20% Bis-Tris gels (Bio-Rad) and visualized by Coomassie blue staining.

### Tubulation Assays

Tubulation reactions containing 2.5 μM retromer complex, 8.75 μM adaptor protein (SNX3, SNX12, SNX12(Δ160-172), SNX12(Δ1-25), or SNX12 H133A/P134A/L135A) and 25 μM of L2(441-460) peptide or its mutants were incubated with 600 μM liposomes overnight at 4 °C in tubulation assay buffer. Aliquots of 4 μL were applied to glow-discharged holey carbon grids (Quantifoil® R2/2, Cu 200 mesh), frontside-blotted at 8°C for 2 s at 90% relative humidity, and plunge-frozen in liquid ethane using a Leica EM P2 plunger. Images were acquired on a JEM-2200FS transmission electron microscope operating at 200 kV and equipped with a K2 direct detector.

### Crystallization and Structure Determination

Crystallization of the VPS35(14-470)-VPS26A-L2(449-458) complex with SNX12 was performed following a protocol similar to that previously described for the Retromer-SNX3 complex ^18^. Briefly, the VPS35(14-470)-VPS26A-L2(449-458) complex (50 µM) was mixed with a three-fold molar excess of SNX12, and the L2(449-458) peptide was included as an additive (150 µM), in buffer containing 300 mM NaCl, 1 mM DTT, 25 mM Tris-HCl (pH 7.5) prior to crystallization trials. Crystals were obtained by mixing 3 µL of protein solution and 1 µL of precipitant solutions containing 0.08 M lithium citrate and 0.88 M ammonium sulphate. Streak seeding was required to obtain diffraction-quality crystals. Crystals were grown by hanging-drop vapor diffusion at 18°C, and oval-shaped crystals appeared within one week. Crystals were flash-frozen in liquid nitrogen using the crystallization solution supplemented with 25% of ethylene glycol as cryoprotector.

Data collection was carried out at ALBA beamline XALOC (Barcelona, Spain) and Diamond Light Source beamline I24 (Didcot, United Kingdom). The structure was solved by molecular replacement using the known structure of VPS26-VPS35(14-470)-SNX3-DMT1-II complex (PDB ID code 5F0J). All diffraction data sets were processed using the XDS program ^37^. The model was built manually in Coot ^38^ and iteratively refined using Phenix ^39^.

### CryoEM grid preparation and data collection

Sample preparation for cryo-ET followed the same procedure as in tubulation assays by incubating 600 µM liposomes with 2.5 μM retromer complex, 8.75 μM SNX12 and 25 μM L2(441-460) in tubulation assay buffer. Before sample vitrification, BSA gold tracers (6 nm; Electron Microscopy Sciences) were added to the tubulation reaction in a 1:4 volume ratio. Four microliters of the sample were frontside-blotted for 2 seconds at a relative humidity of 90% and a temperature of 8°C on a glow-discharged holey carbon grid (Quantifoil® R2/2 Cu 200 mesh grids) before plunge-freezing in liquid ethane (Leica Plunger EM P2). High-resolution cryo-electron tomography data were collected at the cryo-EM facilities of the Electron Bio-Imaging Centre (eBIC, Diamond Light Source, UK) and at the Basque Resource for Electron Microscopy (BREM) at the Instituto Biofisika (UPV/EHU-CSIC) using a Titan Krios transmission electron microscope (Thermo Fisher Scientific) equipped with a BioQuantum energy filter and a K3 direct electron detector (Gatan). The combined datasets comprised a total of 247 tilt series, acquired using the Hagen tilt scheme between -60 and 60 degrees with an angular step of 3 degrees and defocus values between -1.5 µm and -4 µm, in steps of 0.5 µm. For each tilt angle, 10 frames were recorded with a dose rate of 0.3□e^-^ □Å^-2^ per frame and a total tomogram dose of 140□e^-^□Å^-2^. Acquisition magnification was ×53,000, rendering a pixel size of 1.67 Å.

### Preprocessing and tomogram reconstruction

Each movie was motion-corrected and gain-compensated using MotionCor2 ^40^. Each micrograph was Fourier-binned to 5760 x 4092 pixels with a pixel size of 1.67Å. The micrographs were stacked into tilt series using MATLAB and Dynamo ^41^. The tilt series were dose-weighted using a MATLAB script implementing the algorithms introduced in Unblur ^42^. The contrast transfer function was estimated using CTFFIND4 ^43^ and the CTF correction was done using the ctfphaseflip command from IMOD ^44^. A total of 247 tilt series were created. Tilt series alignment and tomogram reconstruction were done using Dynamo Tilt Series Alignment ^45^. After visual inspection, a total of 180 tilt series were deemed viable for reconstruction. The 180 resulting tomograms were reconstructed by weighted back-projection with a pixel size of 6.68 Å and manually annotated.

### Subtomogram averaging

The tomograms were annotated using the Dynamo Catalogue ^46^. A total of 222 retromer-coated tubules were modelled using the filament models by manually selecting approximate points along the centerline of the tubulations. A spline interpolation was done between those points to accommodate the tubulation curvatures. This allowed for the extraction, averaging and alignment of 6864 subvolumes (of 128x128x128 binned pixels) containing sections of the coated tubes. Because the tubules present considerable heterogeneities such as different diameters, this average allowed for the identification of arch complexes only on a section of the tube circumference. However, this was sufficient for the application of an oversampling strategy, generating a set of 80,951 subvolumes (of 64x64x64 binned pixels) in the expected area of the presence of arches along all tubules. The azimuthal angle of all particles was then randomized to remove template biases in following STA project. These 80951 subvolumes were then refined through a second alignment project localized specifically on the retromer (arch) coat. The resulting 57,497 particles generated an average that allowed for the general identification of the local coating architecture. In particular, a two-class MRA project localized on the area beside the central arch was used to identify 19177 particles (of which 16847 were selected for averaging) that presented another arch beside the central one and 15757 particles (of which 15251 were selected for averaging) that did not present such arch.

Finally, the 57,497 particles were locally reconstructed at full scale of 1.67 Å per pixel. The particles were reconstructed to the size of 256x256x256 by applying weighted back-projection to corresponding patches of the tilt series. The patches side-length was three times the particle side-length so as to avoid reconstruction artefacts. The particles were aligned following a gold standard approach on the central arch. The alignment mask was a tight parallelepiped that included the central arch but excluded neighbouring arches as well as the membrane bilayer. The final resolution was estimated at 11 Å by gold standard Fourier shell correlation (FSC) in RELION ^47^. The local resolution map was also computed with RELION.

### Class analysis and model building

As mentioned previously, MRA was used to classify a subset of particles by the presence or lack of a neighbouring arch. The class with the neighbouring arch was named A class (as *A*rch), while the class without the arch was named V class (as *V*alley). As such, these sets of particles were selected to investigate the lateral interactions between retromer chains.

A total of 15,337 particles of size 256x256x256 of class A were re-cropped after centering on the Vps26-Vps26 contact from the neighbouring retromer strand. The particles were aligned with Dynamo and the final resolution was estimated at 12 Å by gold standard FSC in RELION. The alignment mask was a tight parallelepiped that excluded neighbouring retromer chains as well as the inner layer of the membrane.

Finally, a total of 18110 particles of size 128x128x128 of class A were re-cropped after centering on the area between two laterally neighbouring arches. The particles were aligned with Dynamo and the final resolution was estimated at 13 Å by gold standard FSC in RELION. The alignment mask was a parallelepiped that included both neighbouring retromer strands.

Neighbourhood analysis was used to cross-validate results from STA and to build coating models. Neighbourhood analysis consisted of computing the relative positions of pairs of particles. For each particle all other particles closer than a threshold were identified and selected as the neighbours. The distance vectors between the particle and its neighbours were then computed and then rotated by the alignment parameters of the particle. This process was repeated for all particles, creating a statistical description of spatial relationships between arches at short distances.

Neighbourhood analysis of the class A was computed by evaluating the neighbourhood of 15,310 particles assigned to class A to 50,105 particles that were used to generate 11 Å arch density map. A total of 70,960 pairs of particles were included by limiting the maximal distance of particles to be considered neighbours to ∼16 nm. Neighbourhood analysis of the class V was computed by evaluating the neighbourhood of 14,903 particles assigned to class V to the same 50,105 particles. A total of 69995 pairs of particles were included by using the same distance threshold as before.

Additionally, the radius of tubulation was estimated by fitting a bent cylinder model to the positions of all found retromer arches. As the center of each arch subvolume after STA alignment is meant to be on the surface of the membrane bilayer, the STA positions that generate the arch density map of 11 Å of resolution were used as sampled points on the tubulation surface for the purposes of model fitting. A single radius was imposed per tubulation. The results were cross-validated by inspecting the results of PCA analysis in Dynamo on an average of tube sections. The found radiuses were separated into quartiles. The particles found on tubulations with fitted radius in the first quartile (radius smaller than 22 nm) were classified as small radius particles, while the particles in the fourth quartile (radius larger than 27 nm) were classified as big radius particles. This was done to allow for clearer identification of structural differences by inspecting density maps using Chimera UCSF ^48^. Chimera Morph Map tool was used to generate morphing videos.

Neighbourhood analysis of the big radius particles was done by comparing 3,442 particles to the reference 50,105 particles. A total of 21,035 pairs of particles were found. Neighbourhood analysis of the small radius particles was done by comparing 3,919 particles to the reference 50,105 particles. A total of 26,901 pairs of particles were found. These results were used to estimate local distances between arches to parametrize a simplified coating model to explain the co-existence of A and V classes particles in the same coat and discuss the accommodation of different tubulation radiuses.

### Angle and distance estimations

The distances and angles between retromer arches were computed using neighbourhood analysis comparing particles centered on the Vps26-Vps26 contact to particles centered on the surface of the membrane bilayer under an arch. The arch-to-arch distance was estimated by adding the measured Euclidean distances from the VPS26-VPS26 contact to the previous and following arch. The selection of pairs of particles conforming to this geometry was done manually, generating two clusters of pairs of particles indicating the relative positions of arches. The arch-to-arch distance was estimated by fitting a Gaussian distribution to each cluster and selecting the centers. The arch-to-arch distance for big radiuses was estimated to ∼27.9 nm and for small radiuses to ∼27.2 nm. The angle between the direction of tubulation and the arch-to-arch direction was estimated by projecting the arch position on the plane of the VPS26-VPS26 contact and comparing the vector between these points and the Y-plane of the particles. This angle was estimated to ∼10.0° for big radiuses and ∼11.4° for small radiuses.

The VPS26-VPS26 contact hinge-like motion was visualized by comparing the density maps for big and small radiuses of the particles class A that were re-cropped and aligned after centering on the area between two laterally neighbouring arches. The field of view was increased by changing the average side-length from 128 pixels (21 nm) to 256 pixels (42 nm).

The lack of similar arch motion was visualized by comparing the arch densities maps for large and small radiuses, maintaining the 256 pixels (42 nm) sidelength.

### Template matching analysis

Template matching was used on a sub-set of tomograms to visually investigate specific coating arrangements. The GUI functionalities of model-aware template matching in Dynamo was used ^49^. The tomograms with pixel size of 6.68 Å was utilized. The arch density map at 11 Å of resolution was rescaled to 64x64x64 with a pixel size of 6.68 Å and used as a template. The template mask was the same tight parallelepiped mask used during STA. The tomograms were chosen for having a visually mostly straight tubulation. The tubulation area was identified by manually creating a single oblique chunk of sufficient size. The template angles during computation of template matching were restricted to a cone whose axis is collinear with the direction of tubulation and with a cone aperture of 60 degrees. The azimuthal range was kept to 360 degrees. The angular sampling steps were 5 degrees for both cone and azimuth. The resulting cross-correlation maps were also expressed in cylindrical coordinates by manually determining a cylinder axis along the tubulation.

### Idealized model

An idealized model was generated for a tubule with a 25 nm radius, defined by five helical starts, a 27 nm spacing between arches, and a helix angle of 10°. The position of the first arch along each helical start was staggered between even and odd starts to empirically reproduce the lateral offset of arches observed in the A-class density map. A second idealized model was generated for a larger tubule with a 27 nm radius, using six helical starts, an inter-arch spacing of 29 nm, and the same helix angle of 10°.

### Data and code availability

Atomic coordinates and structure factors for the crystallographic complexes have been deposited in the Protein Data Bank (PDB) under the accession code 9Q8M listed in Table S1. Cryo-ET reconstructions and representative tomograms have been deposited in the Electron Microscopy Data Bank (EMDB), with accession codes EMD-56177 and EMD-56426 listed in Table S2. Tilt series and unaligned micrographs have been deposited in the Electron Microscopy Public Image Archive (EMPIAR) under the dataset identifier EMPIAR-13262. Additional data supporting the conclusions of this study are available from the corresponding authors upon reasonable request.

## Supporting information

Five-start helical organization of the SNX12-retromer coat

Six-start helical organization of the SNX12-retromer coat

Hinge flexibility at VPS26-VPS26 junctions

## Acknowledgments

We thank Rocío Alonso from the Hierro Lab and Carlos Fernández-Rodríguez from the Castaño Lab for technical assistance, and Andrés Jimenez from Instituto Biofisika for advanced IT support. This work was supported by the Ministry of Science, Innovation and Universities of Spain (MCIN/AEI/10.13039/501100011033) under projects PID2023-151986NB-I00 (to A.H.), PID2021-127309NB-I00 and PID2024-158469NB-I00 (to D.C-D), by the Swiss National Science Foundation SNF 320030-236069 (to D.C.-D.), and by the United States National Cancer Institute R35CA242462 (to D.D.). This work was supported in part by the Fundación Biofísica Bizkaia and the Basque Excellence Research Centre (BERC) program of the Basque Government. This study made use of the Diamond Light Source (Oxfordshire, UK), ALBA synchrotron beamline BL13-XALOC, the cryo-EM facilities at the UK national Electron Bio-imaging Centre (eBIC), the Basque Resource for Electron Microscopy (BREM) at Biofisika Institute, the X-ray and electron microscopy platforms at CIC bioGUNE, and the Flow Cytometry Core at the Yale Cancer Center.

## Author contributions

M.P.P. performed cloning, purification, crystallization, structure solution, ITCs, co-flotation assays, cryoEM and cryoTEM; R.C. performed cryoTM, subtomogram averaging and structure analysis; C.O. and P.Z. generated and analyzed the knockout and knockdown cell lines; A.L.R. performed X-ray crystal structure solution, cryoEM and cryoTEM. D.D., D.C.D. and A.H. designed the research, analyzed the data and wrote the manuscript.

## Competing interests

The authors declare no competing interests.

## Extended data figures and tables

**Extended Data Fig. 1:**
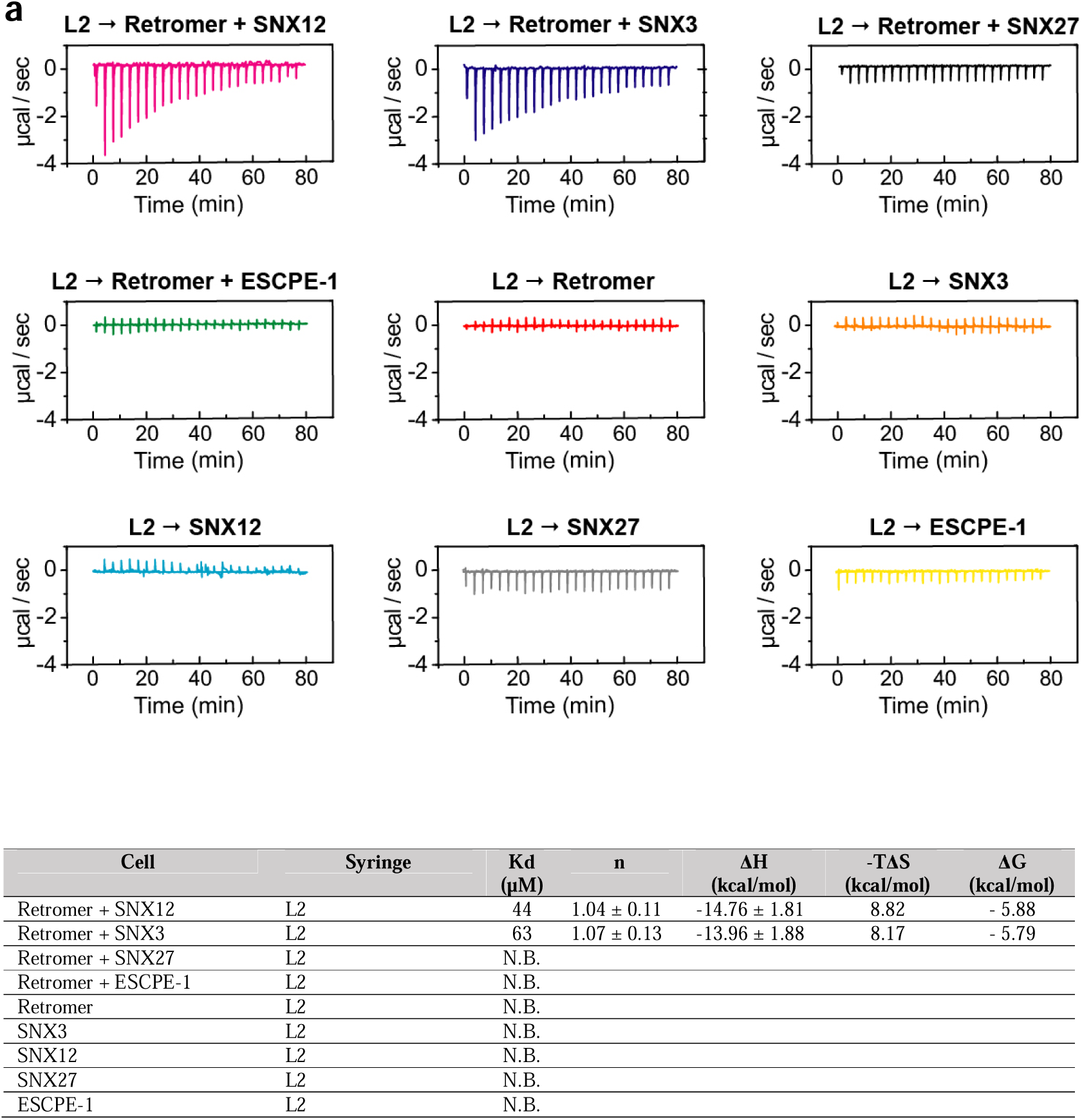
Raw ITC data related to Figure 3a. (**a**) Thermograms obtained from representative ITC titrations showing the interaction of the HPV L2(441-460) tail with retromer and retromer-SNX-associated proteins involved in different endosomal sorting routes. (**b**) Summary of ITC thermodynamic parameters presented in (**a**).

**Extended Data Fig. 2:**
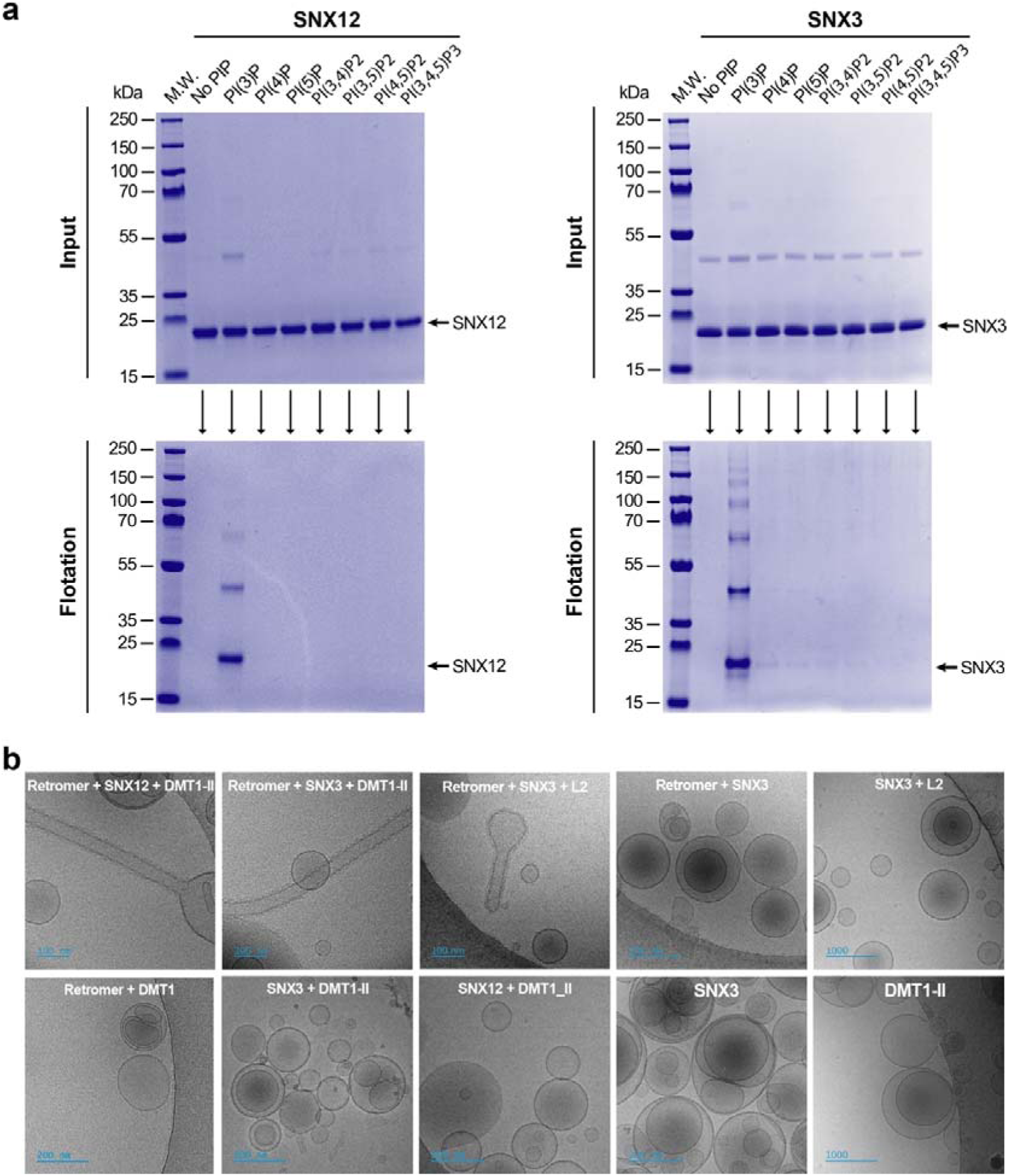
PI3P binding and cargo-dependent membrane tubulation by retromer SNX complexes. (**a**) Liposome flotation analyses showing selective binding of SNX12 (left panel) and SNX3 (right panel) to PI3P among tested phosphoinositides. (**b**) Cryo-EM analysis showing cargo-dependent membrane tubulation. Tubule formation occurs only in the presence of retromer, an SNX adaptor, and the DMT1-II(550-568).

**Extended Data Fig. 3:**
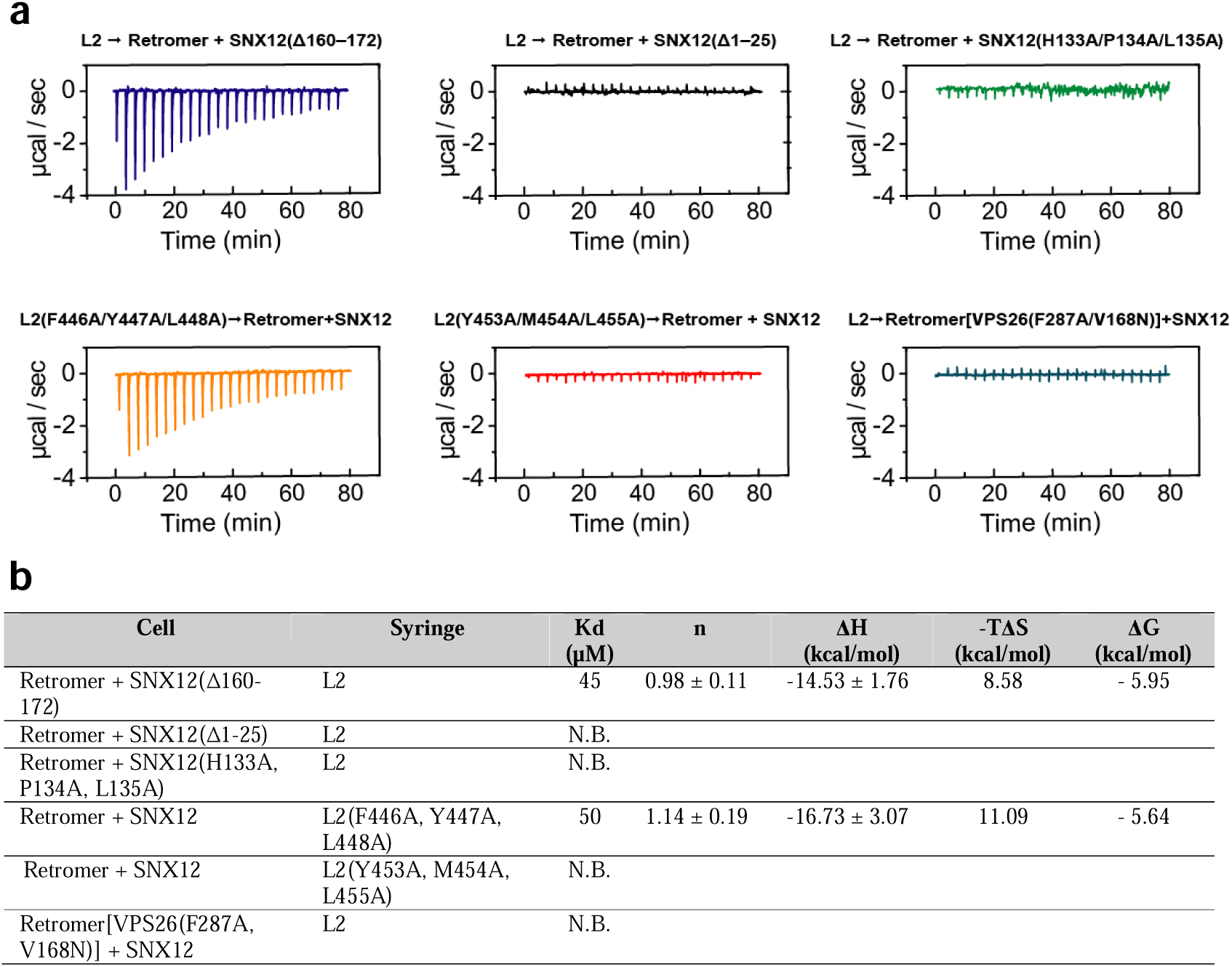
Raw ITC data related to Figure 4d. (**a**) Thermograms obtained from representative ITC titrations showing the interaction of the HPV L2(441-460) tail with retromer-SNX12 complexes and the indicated mutants. Mutations include deletions of the SNX12 N terminus or C terminus and substitutions disrupting the VPS26-SNX12 interface, the VPS26 binding pocket, or the L2 FYL (aa 446-448) and YML (aa 453-455) motifs. (**b**) Summary of ITC thermodynamic parameters presented in (**a**).

**Extended Data Fig. 4:**
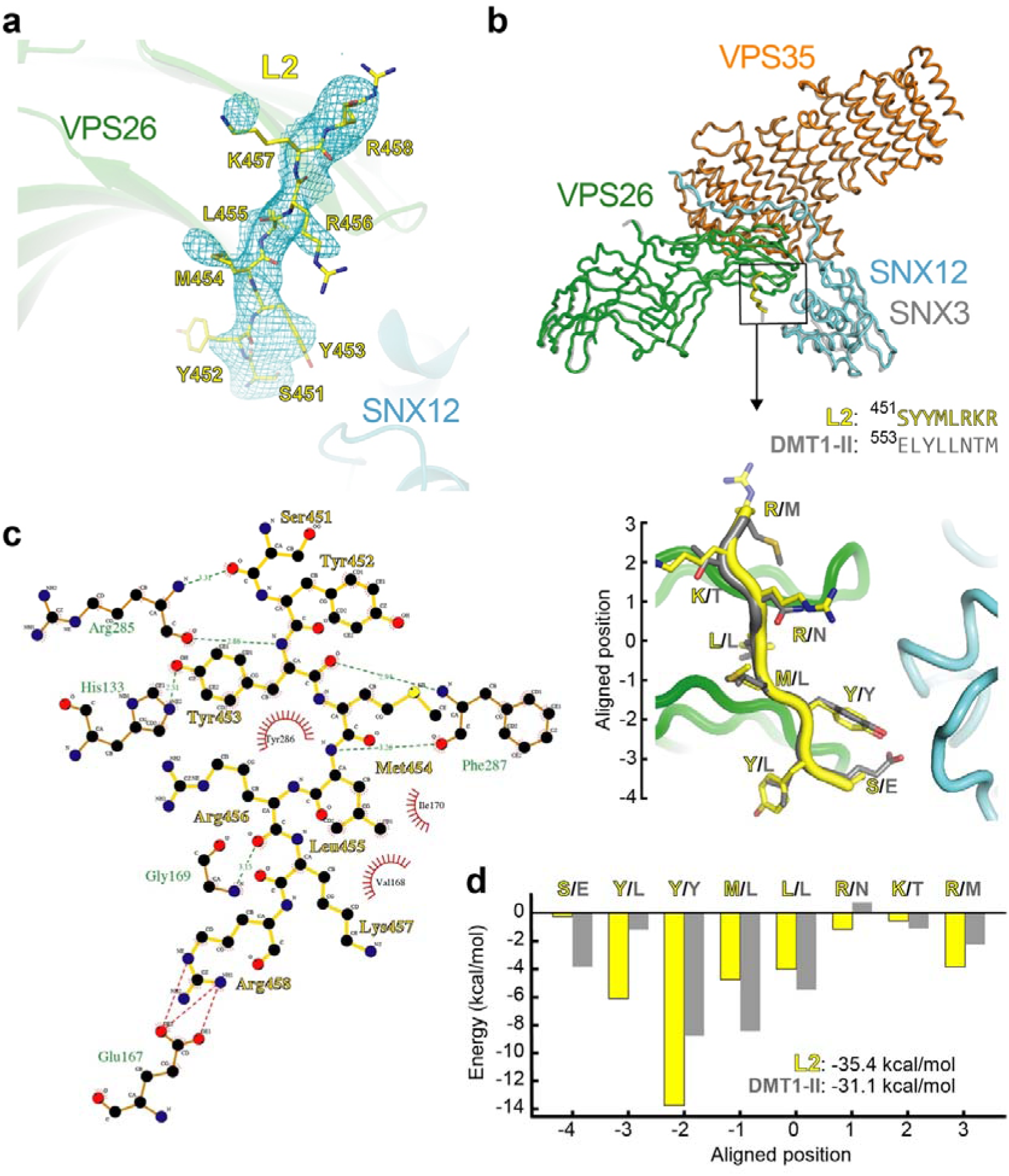
Crystal structure of the Retromer-SNX12-L2 complex reveals conserved cargo recognition features. (**a**) Electron density map (2Fo-Fc) calculated with phases derived from the final refined model and contoured at 1.5σ in the vicinity of the VPS26-VPS35N-SNX12 interface, showing the refined L2 structure in stick representation. (**b**) Structural superposition of the Retromer-SNX12-L2 complex (this work, PDB 9Q8M) and the Retromer-SNX3-DMT1-II complex (PDB 5F0L) shown as tube cartoons, with an inset highlighting the superposed L2 and DMT1 cargo peptides. The alignment, encompassing 906 residues, yields an overall RMSD of 0.387 Å. In PDB 9Q8M (this study), VPS26 is shown in green, VPS35N in orange, SNX12 in blue and L2 in yellow; the corresponding subunits in PDB 5F0L ^18^ are shown in grey with 50% transparency. (**c**) Interaction network at the L2-Retromer-SNX12 complex interface represented in 2D using LigPlot+ ^51^. (**d**) Per-residue energy contributions to the binding affinity of L2 and DMT1 peptides to Retromer-SNX12 and Retromer-SNX3, respectively, estimated using pyDockEneRes ^52^.

**Extended Data Fig. 5:**
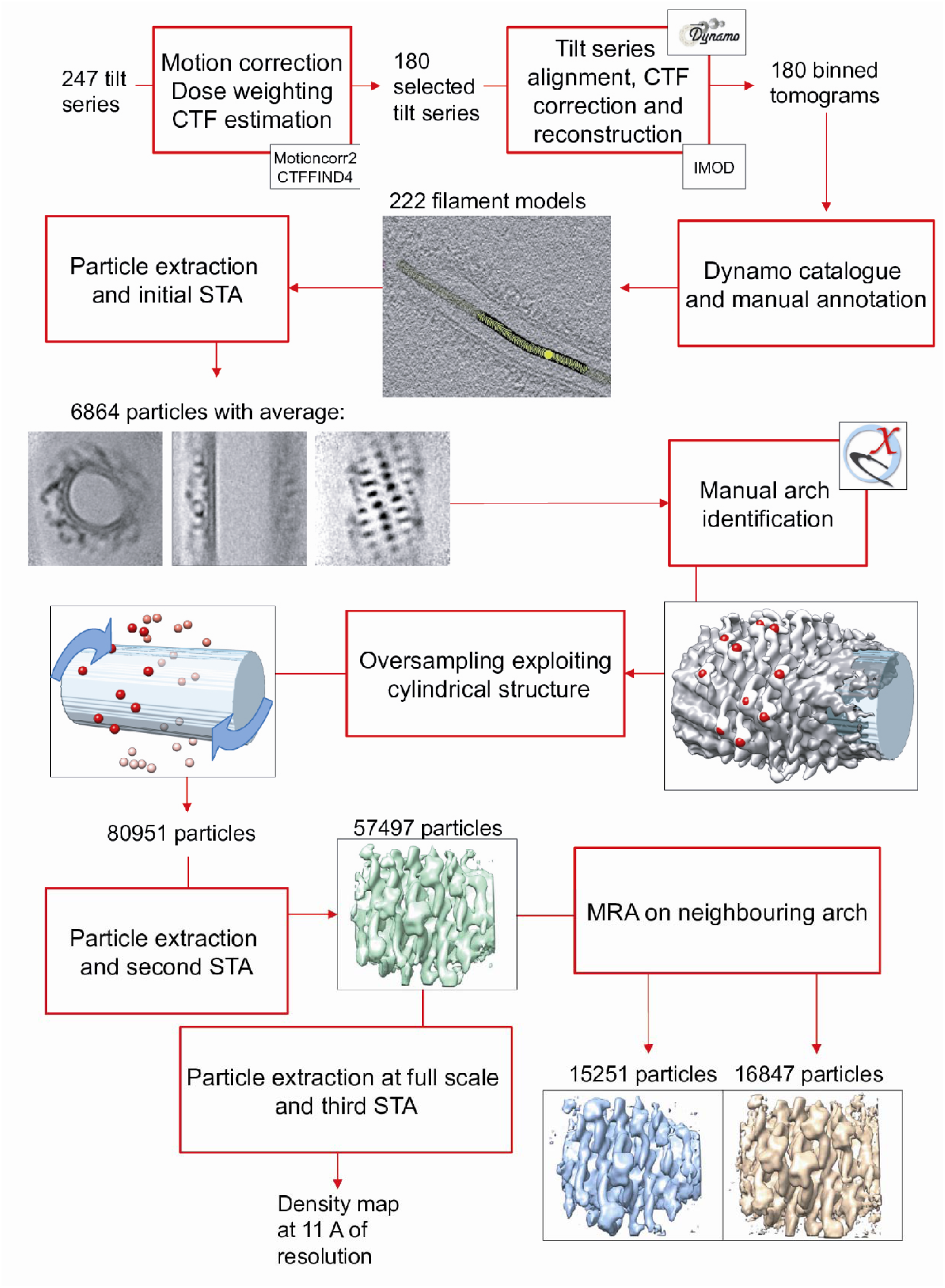
Processing workflow of cryo-ET data. Overview of the processing steps followed to obtain the final average and the two class-specific averages.

**Extended Data Fig. 6:**
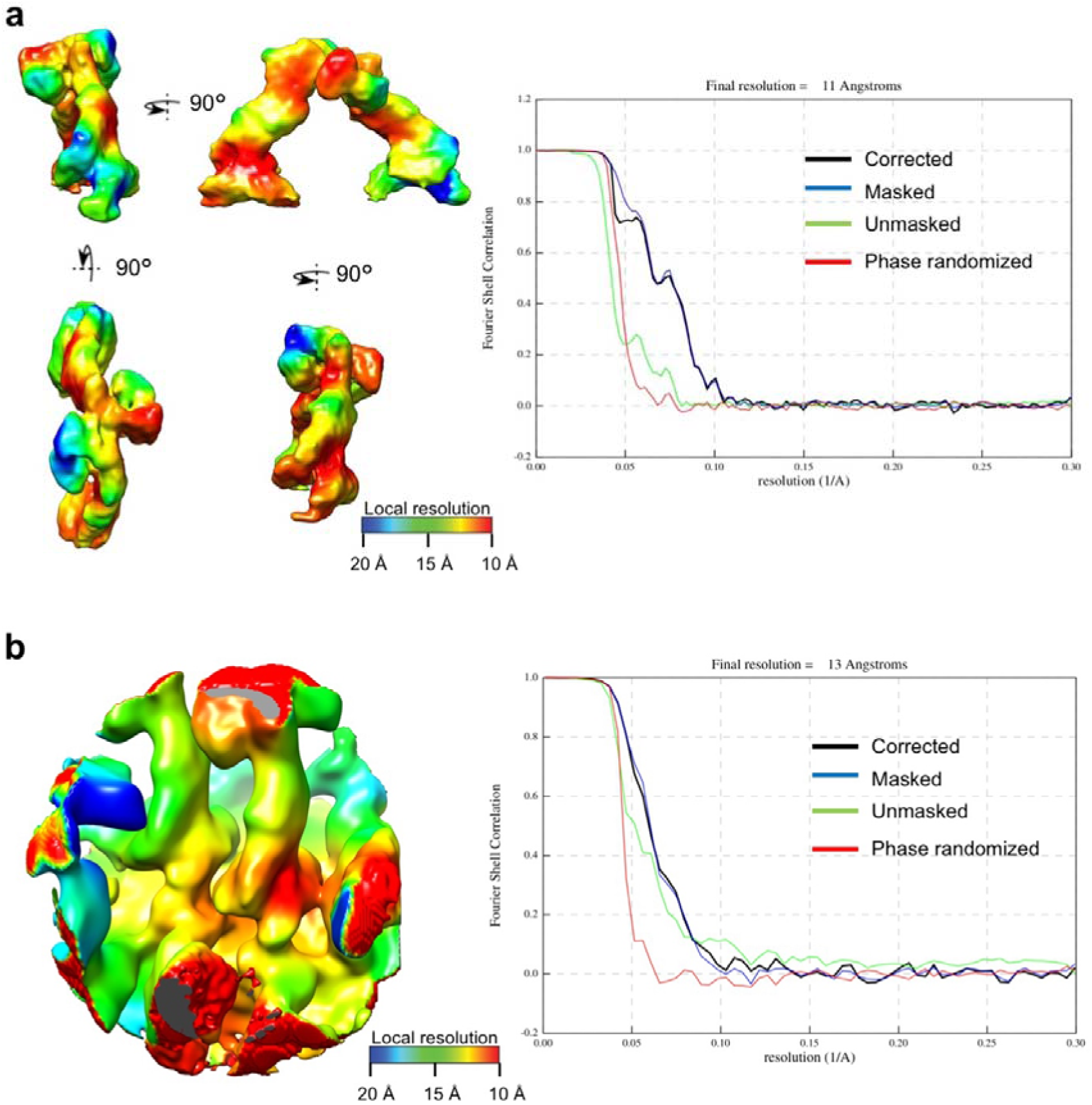
Resolution estimation of the retromer arch and the SNX12-VPS26 lateral contacts. (**a**) Local resolution maps and gold-standard FSC curves showing the estimated global resolution of the arch formed by the retromer-SNX12-L2 complex. (**b**) Local resolution maps and gold-standard FSC curves showing the estimated global resolution of the SNX12-VPS26 lateral contacts.

**Extended Data Fig. 7:**
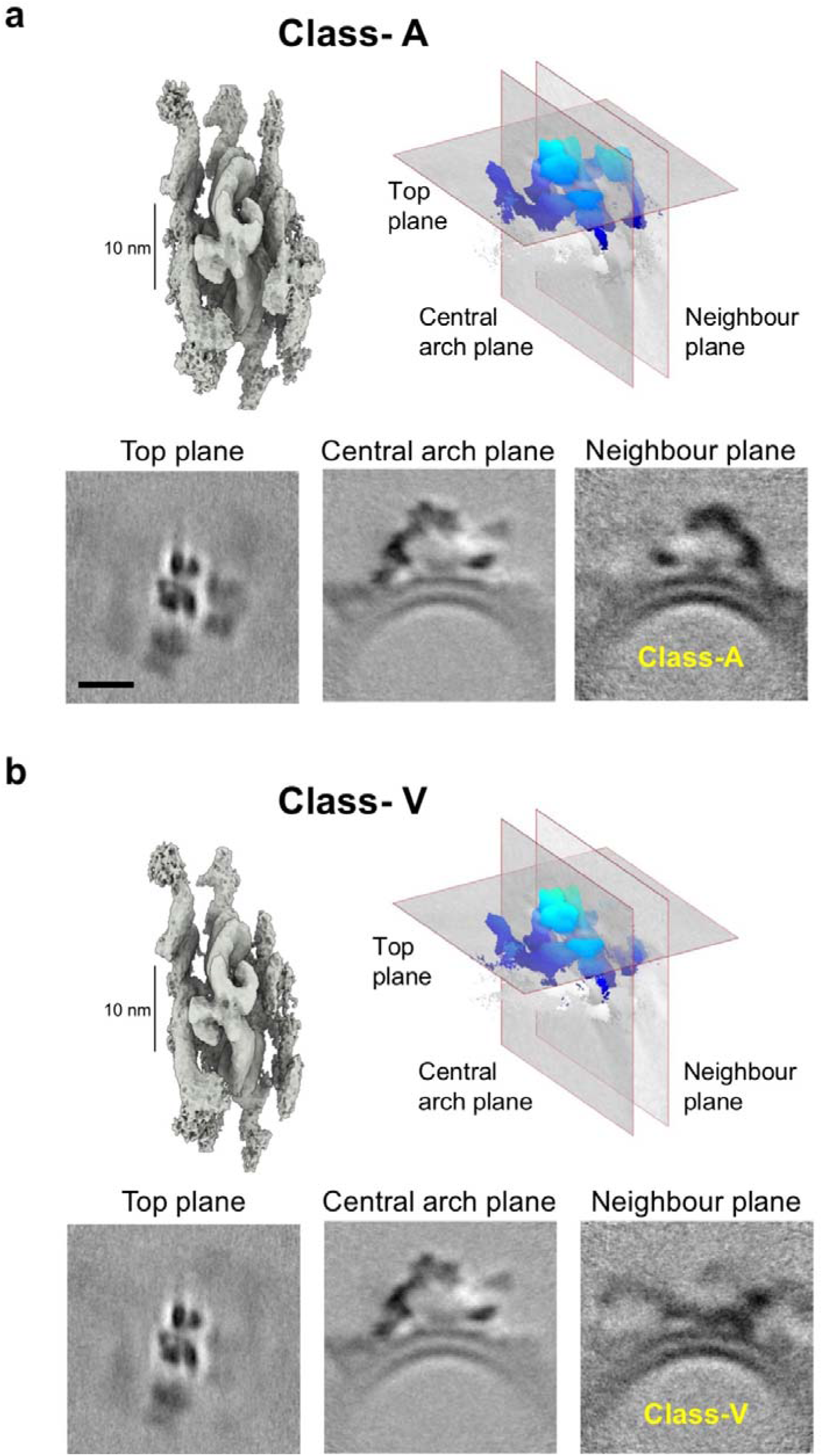
Comparison of class-A and class-V assemblies. (**a**) Global density map of class-A with representative slices illustrating its overall organization. Scale bars, 10 nm. (**b**) Corresponding density map and representative slices of class-V showing the alternative organization.

**Extended Data Fig. 8:**
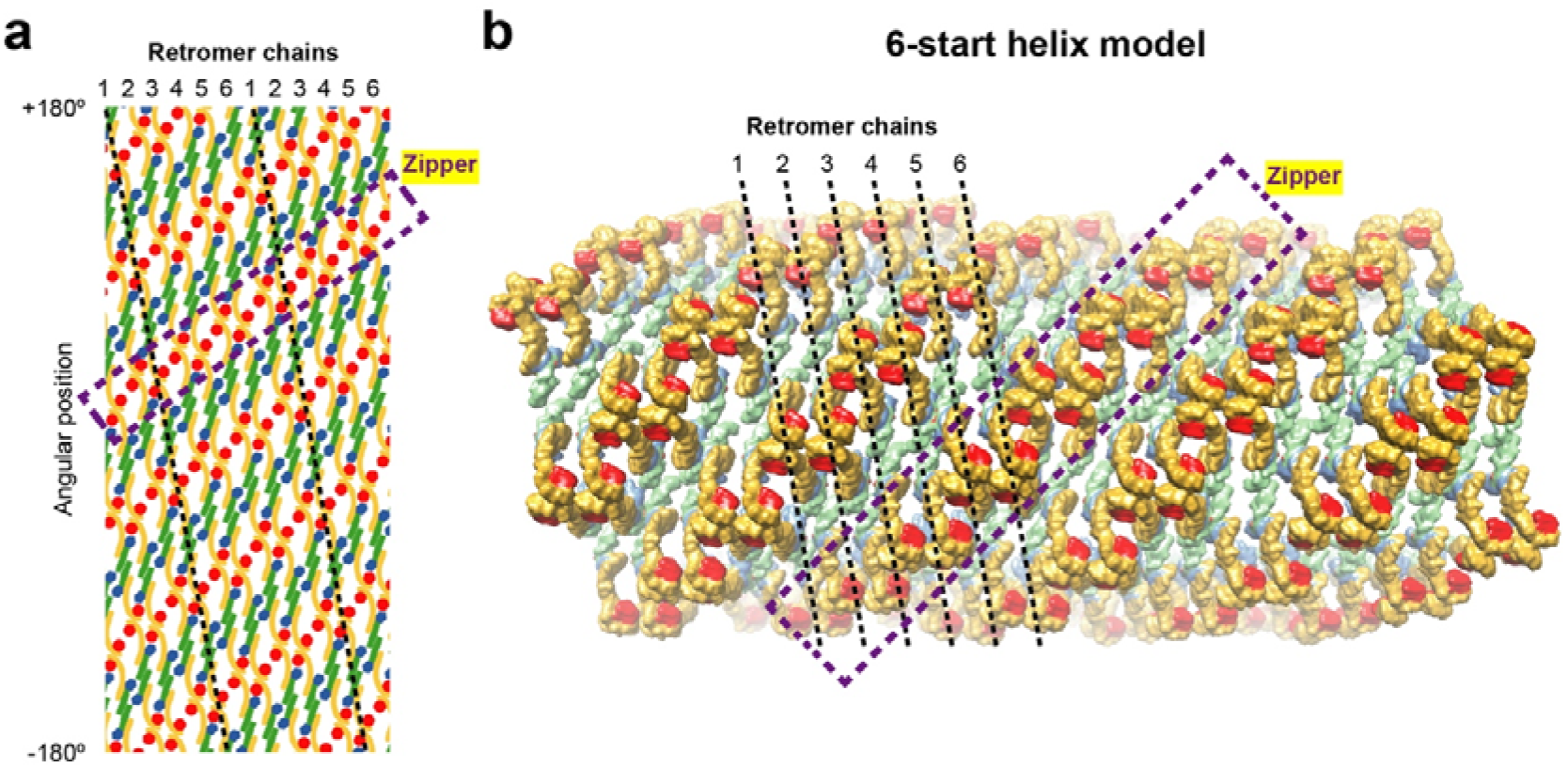
Schematic representations of an idealized 6-start helical model of the Retromer-SNX12 coat. (**a**) 2D schematic representation in cylindrical coordinates. The helical starts are numbered 1 to 6. The helical start numbered 1 is highlighted. (**b**) 3D visualization of the helical arrangement, defined by a 10° chain angle, a tube radius of 29 nm, and an arc-to-arc distance of 26 nm.

**Extended Data Fig. 9:**
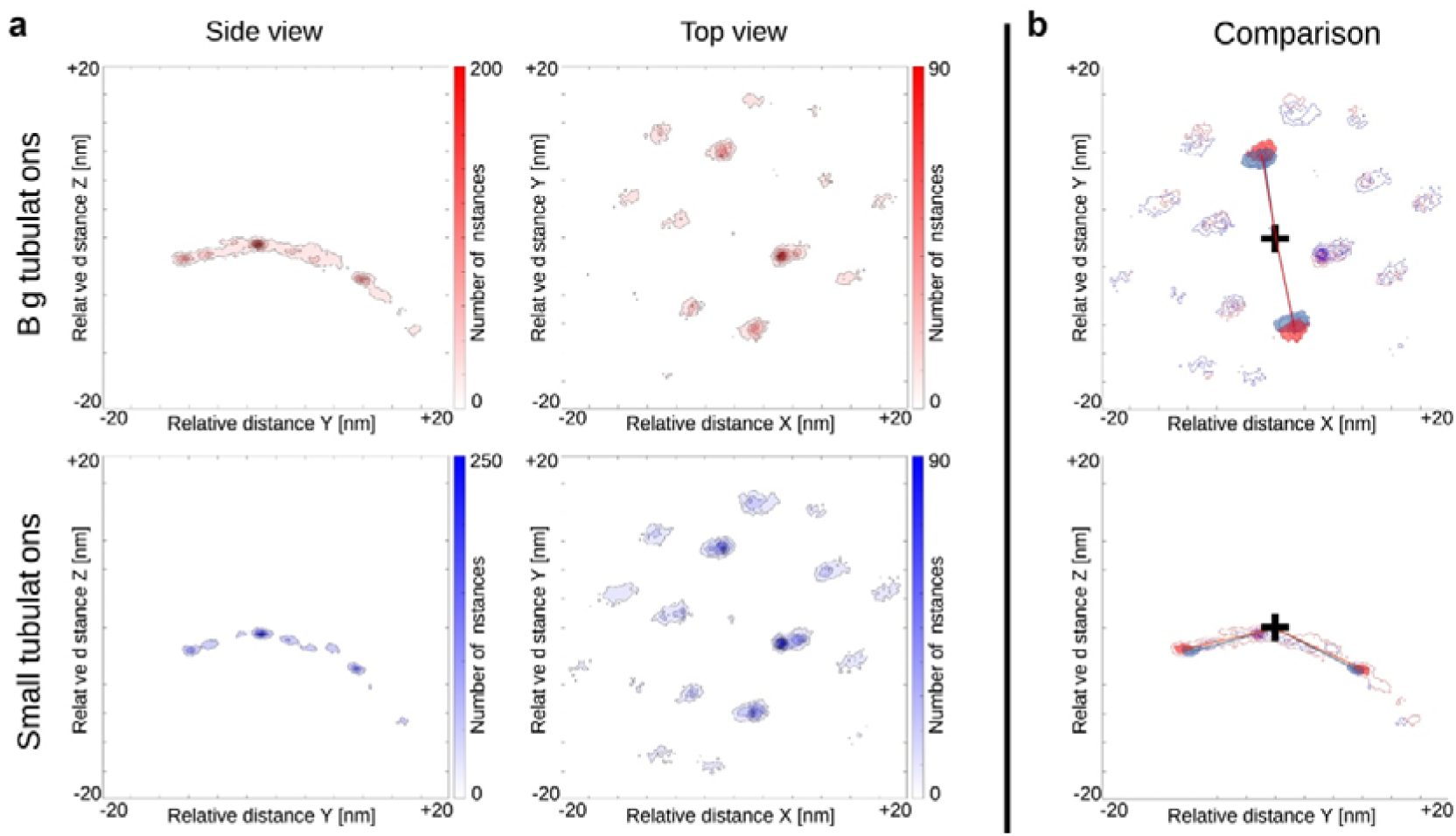
Relative spatial distribution of retromer arches in tubules of different diameters. (**a**) Heatmaps showing the relative spatial positions of retromer arches with respect to the VPS26 dimers used as reference points. Data for tubules with larger radii (more than ∼26 nm) are shown in red (upper panels), and for tubules with smaller radii (less than ∼22 nm) in blue (lower panels). The estimated projections of the retromer arch apices onto the tubule membrane are considered as the arch positions for the purposes of this figure. The x-axis represents the approximate longitudinal axis of the tubule, and the z-axis corresponds to the normal vector of the membrane at the reference sites. Both top and side views are shown. (**b**) Comparison of the relative arch positions between large- and small-diameter tubules. Superimposed heatmaps from both populations are displayed as top (upper panel) and side (lower panel) views. The central position of the reference VPS26 dimers is indicated by a black cross. Regions corresponding to the two retromer arches anchored by the VPS26 dimers are highlighted in red and blue, with connecting lines illustrating the retromer arch-VPS26 dimer-arch relationship and their relative displacement between the two tubule populations.

**Extended Data Table 1:** X-ray crystallography data collection and refinement statistics.

| VPS35<sup>14-470</sup>-VPS26<sup>1-323</sup>-L2<sup>449-458</sup>-SNX12-L2<sup>449-458</sup> |  |
| --- | --- |
| **Data collection** |  |
| Wavelength [Å] | 0.97926 |
| Space group | C121 |
| Resolution [Å] | 91.78-3.13 (3.5-3.1) |
| Cell dimensions: |  |
| $a$, $b$, $c$ [Å] | 370.51, 74.63, 57.72 |
| $\alpha$, $\beta$, $\gamma$ [°] | 90.00, 97.74, 90.00 |
| CC<sub>1/2</sub> (%) | 99.7 (52) |
| Completeness (%) | 86.6 (58.7) |
| I/$\sigma$ | 8.2 (1.7) |
| Number of unique reflexions | 14,019 (701) |
| Redundancy | 6.7 (6,8) |
| **Refinement** |  |
| R-factor (%) | 21.4 |
| R-free (%) | 25.3 |
| Number of atoms: |  |
| Protein | 7477 |
| Water | 7 |
| Ion | 4 |
| SO<sub>4</sub> | 4 |
| B-factors |  |
| Protein | 108.35 |
| Water | 56.80 |
| Ion | 134.48 |
| r.m.s. deviations |  |
| Bond length (Å) | 0.002 |
| Bond angle (°) | 0.45 |
| **PDB ID code** | **9Q8M** |

**Extended Data Table 2:**
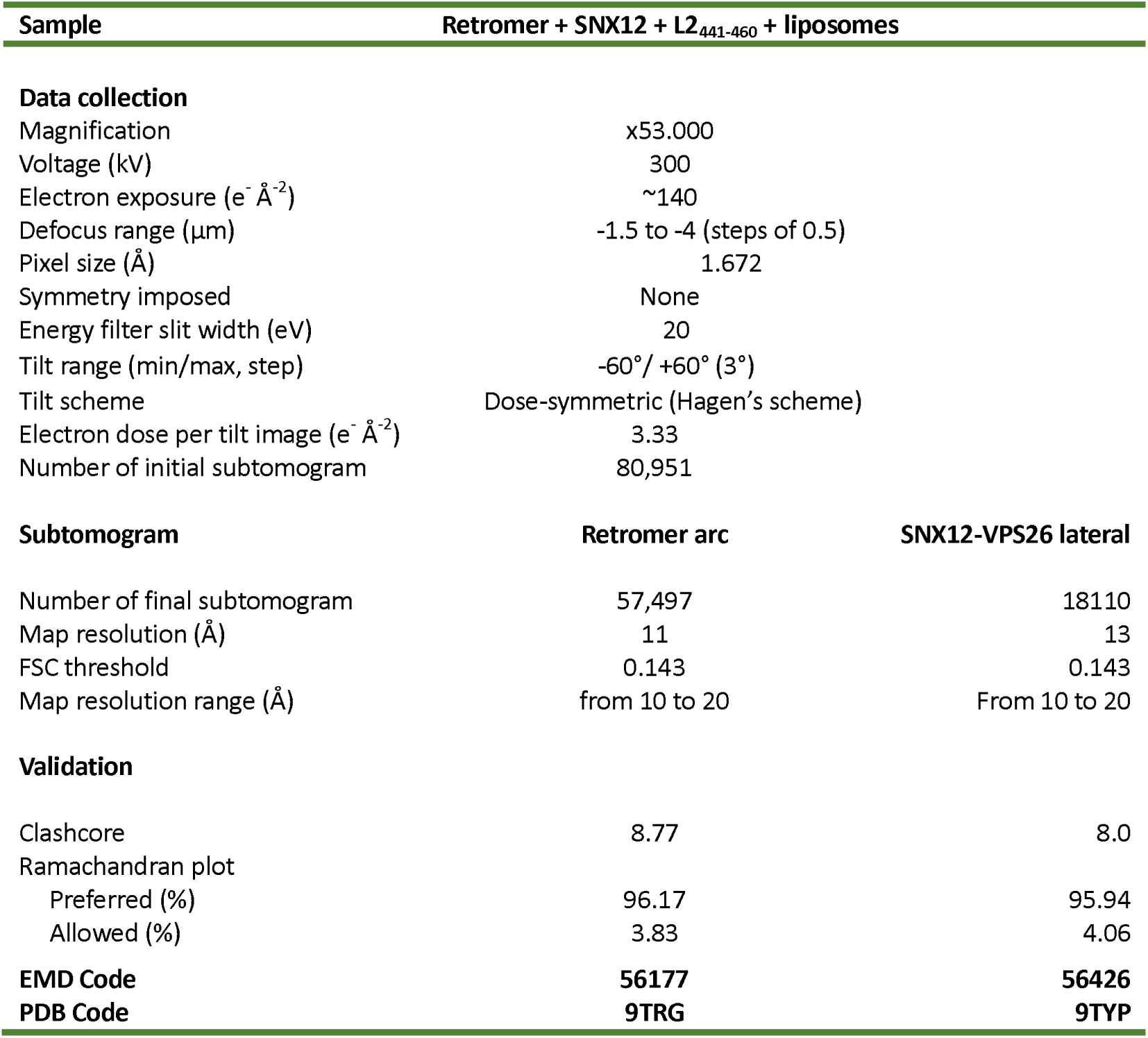
Cryo-EM data collection, processing and model statistics.

**Extended Data Table 3:**
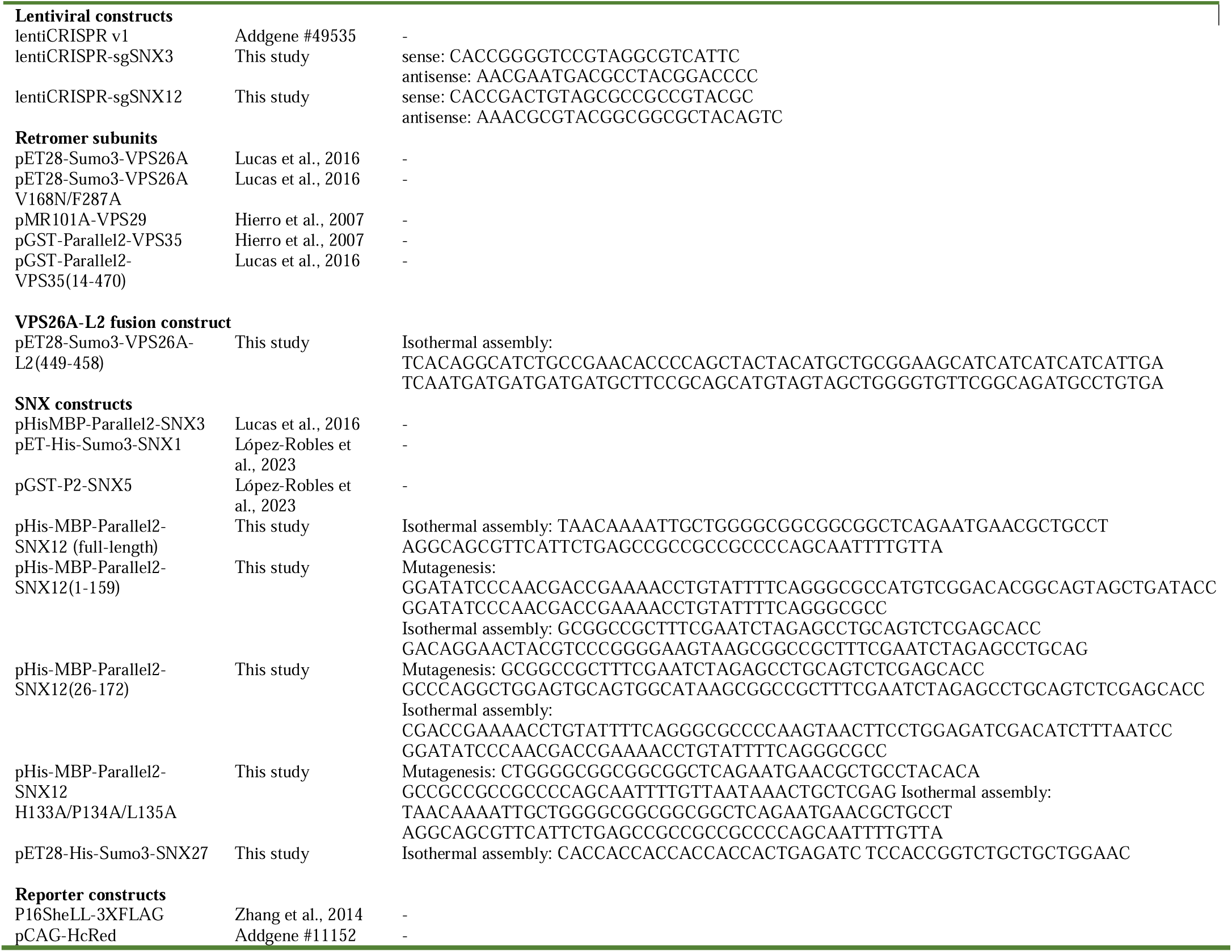
Plasmids and oligonucleotides used in this study.

## Supplementary Information

Supplementary information includes:

- Movie 1: Five-start helical organization of the SNX12-retromer coat.
- Movie 2: Six-start helical organization of the SNX12-retromer coat.
- Movie 3: Hinge flexibility at VPS26-VPS26 junctions enables coat adaptation to membrane curvature.
- Validation report for EMDB ID EMD-56177 and PDB ID 9TRG
- Validation report for EMDB ID EMD-56426 and PDB ID 9TYP
- Validation report for PDB ID 9Q8M

## Notes

### Competing Interest Statement

The authors have declared no competing interest.

